# 3D-Printable and Cytocompatible Hydrogel from *Acinetobacter baylyi* ADP1 Extracellular Matrix

**DOI:** 10.64898/2026.09.11.750904

**Authors:** Shahla Radmehr, Petra Cassiani Ingoni, Ali Eftekhari, Laura Ylä-Outinen, Vijay Singh Parihar, Mohammad Khavani, Noora Perho, Jack Morikka, Dario Greco, Timo Laaksonen, Minna Kellomäki, Heli Skottman, Suvi Santala, Ville Santala

## Abstract

Tissue engineering has advanced significantly, yet multicomponent hydrogels inspired by the compositional complexity of natural extracellular matrices (ECMs) are still underexplored. Most current hydrogels are based on single-component formulations, which can limit their biochemical and mechanical versatility. Developing synthetic multicomponent hydrogels remains challenging because it requires the controlled integration of multiple functional groups within a single material platform. Here, a biologically driven strategy is introduced by leveraging *Acinetobacter baylyi* ADP1, a bacterium that naturally produces extracellular polymeric substances (EPS) composed of a multicomponent matrix of polysaccharides and proteins. Through three-day cultivation and a simple extraction method, a hydrogel is obtained that can be methacrylated and photocrosslinked using red or blue light. This hydrogel is porous, cytocompatible, 3D-bioprintable, injectable, and undergoes rapid gelation for *in situ* crosslinking. This work highlights the potential of using bacterial-derived multicomponent hydrogels for biofabrication.

## Introduction

Tissue engineering (TE) has become an important approach in biomedical applications, aiming to restore or replace damaged tissues and organs through the integration of cells, biomaterials, and bioactive cues^1^. A key requirement is engineered extracellular matrices that provide structural support and direct cellular behavior to promote regeneration^2^. Despite advances in cell biology and biofabrication, developing suitable scaffolds for diverse tissues remains an ongoing challenge^3^.

Hydrogels are widely used in different biomedical applications and biofabrication including TE, because of their water content, tunability, and cytocompatibility^4^, yet both natural and synthetic systems have limitations. Natural polymers such as collagen and gelatin require complex purification and may raise immunogenic or ethical concerns, whereas synthetic polymers such as polyethylene glycol often lack inherent bioactivity and have limited mechanical tunability^5,6^. Additionally, most hydrogels are single-component systems, which can limit their functional versatility for tissue engineering, including their ability to provide multiple biochemical cues, tunable mechanics, and structural features within one material.

Although multicomponent hydrogels could improve tissue mimicry, their synthetic production is often challenging due to complex chemistries, purification demands, and scalability constraints^7–9^. In parallel, 3D-bioprinting enables precise spatial organization of cells and biomaterials^10^, but many existing hydrogels remain suboptimal as bioinks, motivating new approaches for multicomponent materials for biofabrication.

Bacterial extracellular polymeric substances (EPS) are promising scaffolding materials for TE. EPS are naturally secreted high-molecular-weight biopolymers composed of polysaccharides, proteins, and nucleic acids,^11^ and their variable composition and molecular weight give rise to high structural diversity^12^. In bacteria, EPS support biofilm formation and environmental resilience^13^, and their production is strongly influenced by culture conditions such as nutrient availability, pH, and temperature^14^.

Several bacterial species, such as *Komagataeibacter*, *Pseudomonas*, and *Streptococcus*, are already used to produce medically relevant biopolymers and hydrogels like cellulose, alginate, and hyaluronate^15,16^. However, most studies focus on homomeric components with limited functionality or intracellular polymers that require cell lysis and purification, largely overlooking the potential of multicomponent EPS. Key gaps remain in exploring the potential of multicomponent EPS as biomaterials for biofabrication applications and in understanding how environmental and genetic factors can be used to tune their composition and properties.

In this study, we present a novel strategy that utilizes *Acinetobacter baylyi* ADP1 (later referred to as ADP1) as a genetically tractable, non-pathogenic soil bacterium for EPS production. While ADP1 has been studied for its metabolic versatility and transformation capabilities^17^, its potential as a cell factory for EPS-based biomaterials has not yet been explored. Here, we exploit ADP1’s natural ability to produce multicomponent EPS matrices including diverse proteins and polysaccharides. We subsequently methacrylate the EPS and crosslink it with blue and red light to form a cytocompatibility, and highly porous hydrogel. This approach not only circumvents the limitations of single-component systems but also provides a biological route to engineer next-generation biomaterials for 3D-bioprinting and TE.

### Results and discussion EPS production by ADP1

Chemical analysis of ADP1-derived EPS showed ∼75% polysaccharides, ∼20% proteins, and ∼1% DNA, with glucuronic acid, glucose, and rhamnose as the dominant monosaccharides. Such hexose and deoxy-sugar–rich polymers are known to provide rigidity and surface charge, strengthening EPS architecture^18^. Proteomic analysis identified both intracellular (from partial lysis) and extracellular/membrane proteins (Supplementary Table S1), with the most abundant being the biofilm-associated protein (Bap), previously linked to biofilm formation^19^. Bap carried glycosylation motifs (HexNAc and Hex–HexNAc–NeuGc), suggesting a possible glycoprotein role that may contribute to interactions between protein and polysaccharide components within the matrix. To explore the impact of Bap glycosylation, we performed atomistic MD simulations with/without glycans (Fig.1a and supplementary Note 1). Over 150 ns, covalently glycosylated Bap adopted a more defined conformation and showed increased stability based on root mean square deviation (RMSD) trends (Fig. 1b). In simulations where unglycosylated Bap was surrounded by free glycans, both RMSD and gyration (Rg) generally decreased with increasing glycan concentration, indicating a more compact structure, with the lowest Rg at 0.06 M (Fig. 1c, d). Other proteins, including outer membrane protein A (OmpA), tetratricopeptide repeat family protein (TPR), and a hemagglutinin-repeat adhesin (HRA), may also contribute to matrix organization through non-covalent bonds or protein-protein interactions. TPR-family proteins are known to mediate modular protein–protein interactions^20^.

**Fig. 1).**
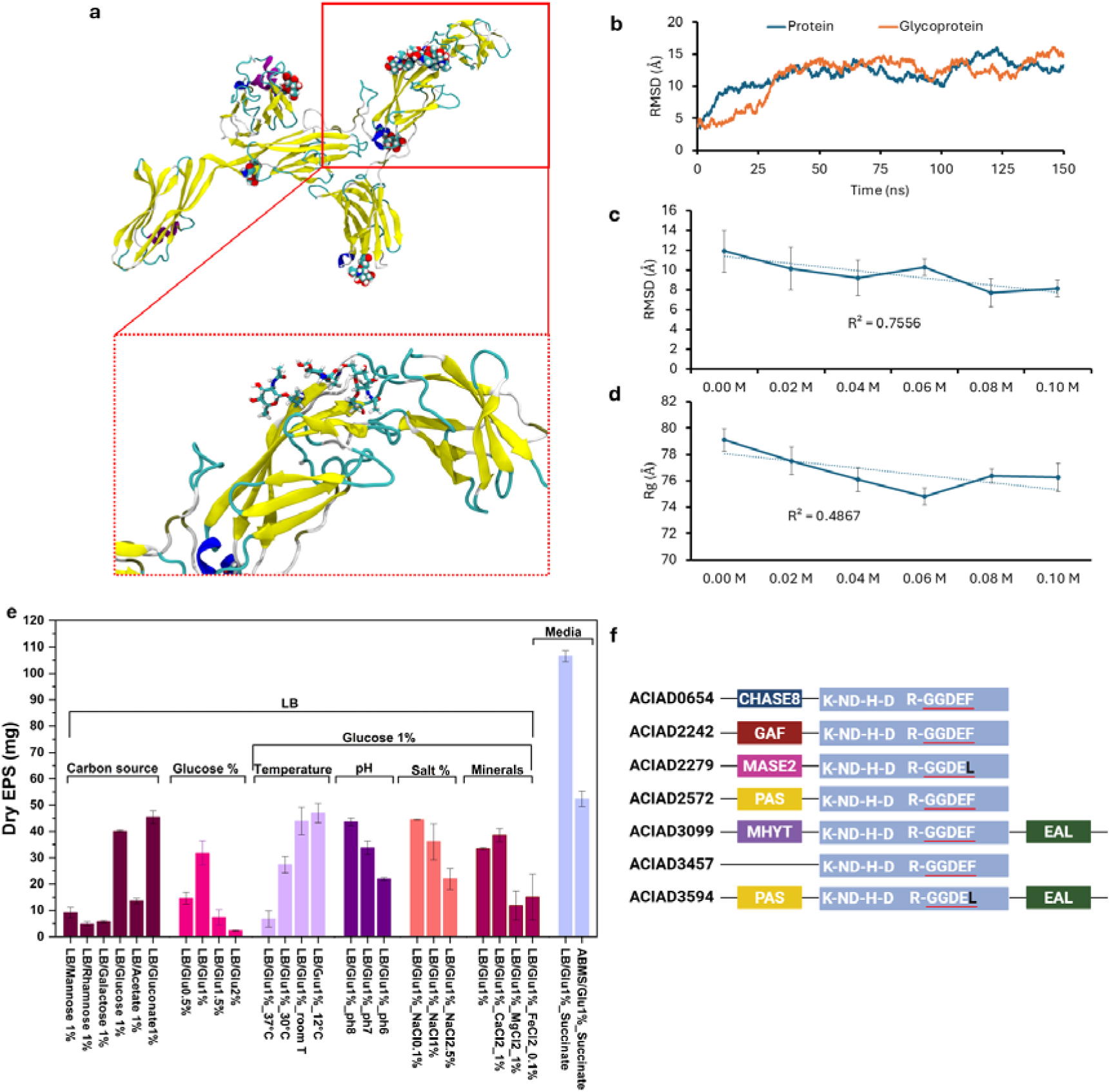
EPS production by *Acinetobacter baylyi* ADP1. **a**) The final structure of the glycoprotein, which can strengthen matrix cohesion by promoting protein–polysaccharide coupling, obtained after 150 ns of fully atomistic MD simulations. **b**) The calculated root mean square deviation (RMSD) values of the protein in the presence (glycoprotein) and in the absence of glycan groups during the simulation time. **c**) The calculated RMSD values of the protein in the presence and absence of glycan groups. In these simulations, in contrast to panels a and b, the protein itself is not covalently functionalized with glycan groups; instead, its dynamical behavior was studied in the presence of different concentrations of freely distributed glycan groups. **d**) The calculated radius of gyration (Rg) values of the protein in the presence of different concentrations of glycan groups. **e**) EPS production is yielded by ADP1 under different culture conditions **f**) Predicted domains of c-di-GMP producing genes.

Given that environmental conditions affect bacterial EPS production^21^, a range of conditions were evaluated (Fig. 1e), starting with different carbon sources. Glucose, gluconate, and acetate were tested as they can support ADP1 growth^22^, while sugars not utilized by ADP1 but detected in EPS (rhamnose/mannose), were tested to assess whether their presence influences EPS production. Glucose and gluconate produced the highest EPS yields (p < 0.05 vs. others) among the tested carbon sources, likely due to their efficient catabolism via the Entner–Doudoroff pathway, which supplies key precursors and reducing power for anabolic biosynthesis. In contrast, in the presence of rhamnose and mannose, the amount of produced EPS was limited. Maximal EPS production was achieved at 1% glucose (p < 0.05 vs. others). Temperature had a significant inverse correlation with EPS production. Among the four tested temperatures (12 °C, room temperature (20-23 °C), 30 °C, and 37 °C), the highest yield was obtained at 12 °C (p < 0.05 vs. 30 °C and 37 °C). This is consistent with studies in *A. baumannii* where low temperatures trigger envelope stress responses which are linked to increased biofilm formation ^23^. EPS production was evaluated across pH values of 6.0, 7.0, and 8.0, revealing a consistent increase with rising pH (p < 0.05 pH 8 vs. others).

Additionally, we tested the impact of divalent cations (CaCl , MgCl , and FeCl ). While calcium had minimal effect, both magnesium and iron significantly reduced EPS production (p < 0.05). Mg may suppress EPS production by stabilizing the outer membrane, alleviating envelope stress, and reducing the need for EPS, as reported for *P. aeruginosa* biofilms^24^. Iron likely acts via the ferric uptake regulator (Fur), which represses stress-related pathways, including those linked to EPS biosynthesis, when intracellular iron is sufficient. Additionally, higher NaCl concentrations (1–2.5%) reduced EPS production. Finally, four media were compared, including LB + glucose, ABMS with succinate as the sole carbon source, LB + glucose + succinate, and ABMS + glucose. The highest EPS yield (112 mg dry EPS) was obtained with LB+ glucose + succinate which is significantly higher than LB + glucose (p < 0.05). One possible explanation is that part of ADP1 growth may be supported by succinate, which could allow some of the available glucose to be redirected toward EPS formation. Fig. S1 shows the images of wet and freeze-dried EPS obtained from cultivations in LB+ glucose + succinate medium.

Previous studies have shown that cyclic bis-(3’,5’)-dimeric-guanosine-monophosphate (c-di- GMP) plays an important role in biofilm formation and exopolysaccharide regulation in bacteria^25,26^, we tested whether it regulates EPS production in ADP1. We identified seven genes in ADP1 carrying GGDEF(L) domains (responsible for c-di-GMP synthesis), with sensory modules (CHASE8, GAF, MASE2, PAS, MHYT, EAL) (Fig. 1f), suggesting possible roles in integrating environmental signals into c-di-GMP pools. Deleting six GGDEF(L) genes (strain ASA1416) did not significantly affect bulk EPS yield compared with wild type under identical conditions (p > 0.05; Supplementary Note 2, and Fig. S2–S5). These findings suggest that the deleted genes do not substantially alter overall EPS production yield under tested conditions.

### Compositional and physical characterization of EPS

Fourier transform infrared spectroscopy (FTIR) results (Fig. 2a) displayed carbohydrate peaks at the 1000–1150 cm ¹ region together with amide bands at ∼1650 and 1540 cm ¹, confirming the multicomponent nature of the matrix. Consistent with prior reviews, microbial biopolymers used in bioprinting (e.g., gellan or hyaluronic acid) are typically purified polysaccharides as rheology modifiers^27^. In contrast, EPS is intrinsically multicomponent, containing both polysaccharides and different protein fractions which places it within the broader category of mixed biological materials such as decellularized-ECM^28^. Decellularized-ECM hydrogels provide complex cues but require tissue sourcing and decellularization. Bacterial EPS production offers an alternative route to multicomponent matrices, with the possibility to tune material yield and composition through cultivation conditions and genetic modification of the production strain. Interestingly, supplementing cultures with different monosaccharides preserved the same functional groups but changed their intensities, indicating a conserved backbone with composition tunable by growth conditions.

**Fig. 2).**
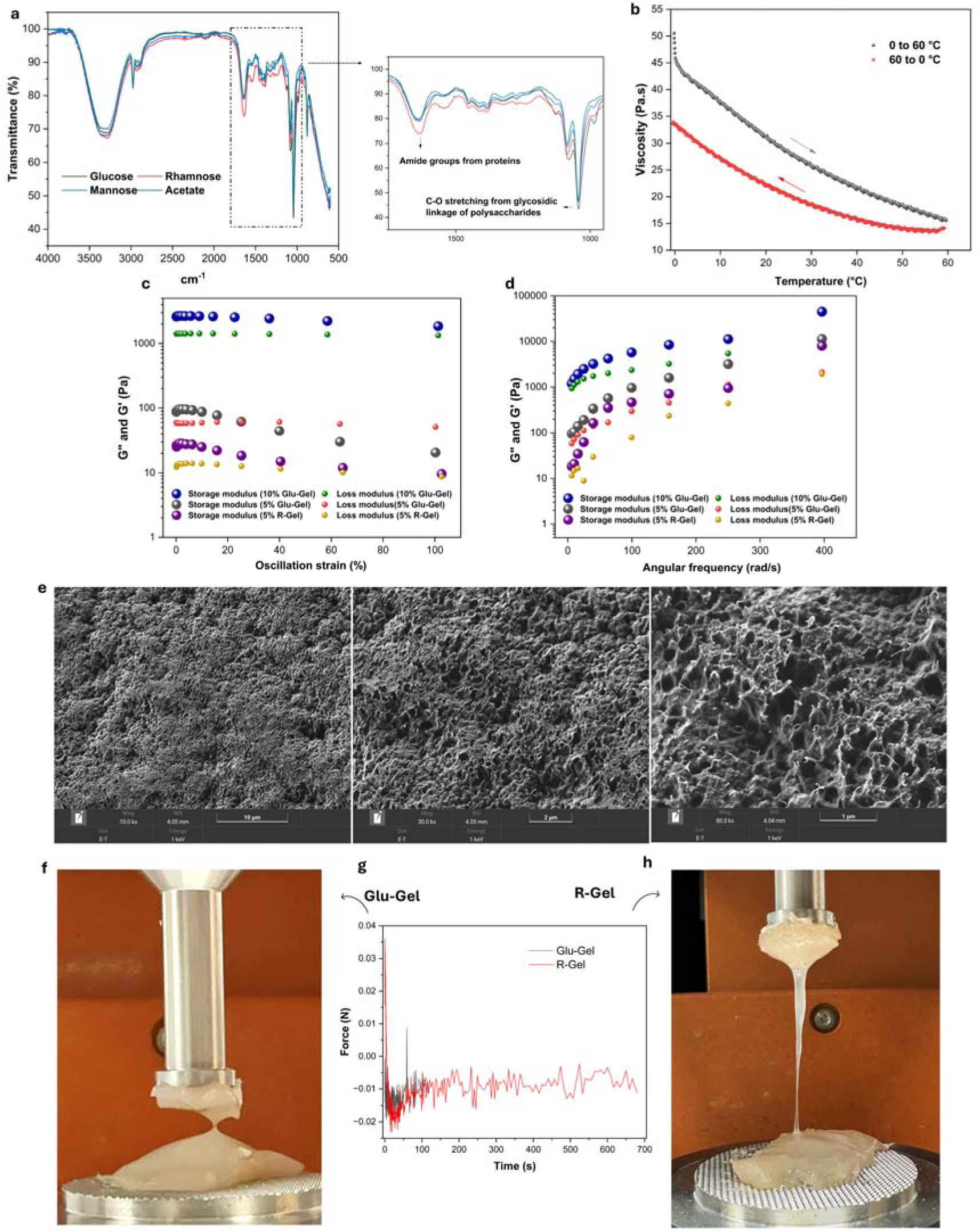
Chemical and physical characterization of EPS-hydrogel. **a**) FTIR spectra showing the presence of protein and polysaccharide in EPS-hydrogel. **b**) Viscosity of EPS-hydrogel while the temperature increased from 0 °C to 60 °C (gray line) and then decreased from 60 °C to 0 °C (red line). **c**) Storage and loss modulus under various oscillation strains. **d**) Storage and loss modulus under various angular frequencies. **e**) SEM images of freeze-dried EPS hydrogel showing a porous microstructure. ImageJ analysis of three independent samples, with 15 measurements per sample, gave an average pore size of 288 ± 51 nm. **f**) Adhesiveness of glucose-based EPS on chicken tissue. **g**) Measurement of tissue adhesion force of hydrogels by the tack adhesion test. **h**) Adhesiveness of rhamnose-based EPS on chicken tissue.

Regarding the physical properties, temperature-dependent viscosity measurements revealed that the EPS hydrogel is influenced by thermal conditions. For a 5% hydrogel, viscosity decreased gradually as the temperature increased from 0 °C to 60 °C, reflecting the weakening of hydrogen bonds and other non-covalent interactions within the protein–polysaccharide network (Fig. 2b, gray). Importantly, when the same sample was cooled back from 60 °C to 0 °C, a substantial portion of the viscosity recovered (Fig. 2b, red), showing that the network is partially reversible. This indicates that the EPS hydrogel can adapt dynamically to temperature fluctuations by breaking and reforming non-covalent links, which can be advantageous compared with widely used thermo-reversible gelatin hydrogels that can soften near physiological temperature^29^. Moreover, the rheological properties of EPS-based hydrogels from ADP1 cultivated on different carbon sources (glucose vs. rhamnose) were evaluated using strain and frequency sweep experiments. Rhamnose was included as a non-metabolizable sugar control, since ADP1 showed similar growth and EPS behavior on LA-rhamnose and LA without added sugar. This comparison was used to distinguish glucose-associated EPS production from the general effect of adding a sugar to the culture medium. In the strain sweep experiments (Fig. 2c), EPS gels produced from ADP1 cultivated on LA-glucose and LA-rhamnose media showed gel-like behavior at low strain, where the storage modulus (G′) remained above the loss modulus (G″). Such a balance between elasticity and viscosity is essential for extrusion-based 3D-printing, where the bioink must flow under shear stress but recover rapidly to maintain the printed shape^10^. The magnitude of G′ scaled with gel concentration, as expected, with the 10% EPS from LA-glucose exhibiting the highest stiffness. Interestingly, the 5% EPS gel produced from ADP1 cultivated on LA-rhamnose showed lower absolute G′ values than the 5% EPS gel produced on LA-glucose, but its mechanical response suggested a more stable elastic regime before network breakdown. The frequency sweep experiments further confirmed these trends (Fig. 2d). In all gels, G′ dominated over G″ throughout the frequency range, indicating a stable elastic network. Notably, many widely used systems improve printability through multi-component formulation by blending components to balance viscosity, stress, and recovery^3^, whereas here these properties emerge from a bacterial-assembled multicomponent EPS matrix. The scanning electron microscopy (SEM) revealed porous and fibrous architecture (Fig. 2e). Such morphology is generally desirable for scaffold materials, as interconnected pores can support fluid transport and cell infiltration ^30^.

A tack test using wet chicken breast tissue was conducted to assess tissue adhesion of EPS-based hydrogels. Both EPS hydrogels produced from ADP1 cultivated on LA-glucose and LA-rhamnose showed measurable adhesion and formed visible bridges during tensile separation. While the absolute detachment forces were relatively low (Fig. 2g), distinct differences were observed in their detachment behavior. The EPS hydrogel produced on LA-glucose failed rapidly with a clean interface detachment (Fig. 2f), whereas the EPS hydrogel produced on LA-rhamnose maintained contact longer, showing filamentous stretching and a more gradual loss of contact (Fig. 2h). Similar results were observed in tack tests without tissue (Supplementary Fig. S6). The gel produced from LB with no added carbon source showed similar behavior to the EPS hydrogel produced on LA-rhamnose (data not shown). The observed wet adhesion may arise from noncovalent polysaccharide–protein interactions between hydroxyl-rich EPS polysaccharides and tissue surface residues. Like catechol- or quinone-mediated hydrogels that adhere via nucleophilic coupling with tissue amines or thiols^31^, the EPS hydrogel may rely on polysaccharide–protein interactions to achieve wet tissue adhesion. Unlike catechol adhesives that require chemical functionalization^32^, the EPS hydrogel shows wet adhesion without catechol/quinone-motifs, suggesting that native interactions support wet adhesion. Such mechanisms are advantageous in physiological environments where maintaining contact and cohesion is critical for stable hydrogel–tissue interfaces. This work provides an early example of using a bacterial multicomponent EPS matrix as a directly printable and injectable hydrogel platform, offering proof of concept potential for biofabrication without the need to construct a synthetic multipolymer formulation.

### Crosslinking and immunotoxicity behavior of EPSMA

To enable light-triggered gelation, methacrylate groups were introduced into EPS to produce methacrylated EPS (EPSMA) (Supplementary Note 3, Fig. S7,8).

Later, crosslinking of EPSMA was achieved by photopolymerization through two distinct mechanisms. Type I reactions rely on photoinitiators that cleave under high-energy blue light, offering rapid initiation but limited applicability^33^. Type II reactions use a photocatalyst combined with electron donors/acceptors, generating radicals at wavelengths ≥ 500 nm^34^.

To demonstrate versatility, *in situ* photorheology was conducted under blue and red light using EPSMA at concentrations of 2.5%, 5%, and 7.5% (w/v), monitoring viscoelasticity by time-sweep rheology (Fig. 3a, b). Under blue light (405 nm) with Lithium phenyl-2,4,6-trimethylbenzoylphosphinate (LAP)^35^, 7.5% and 5% hydrogels rapidly crosslinked, with G′ stabilizing at ∼3.8 and ∼0.9 kPa within 10 min, whereas 2.5% reached only ∼0.1 kPa, indicating insufficient crosslink density. Under red light (625 nm) using methylene blue (MB ) and triethanolamine (TEA)^36^, increasing concentration from 2.5% to 7.5% raised the final G′ from ∼0.8 to ∼8 kPa within ∼15 minutes, reflecting higher crosslink density and chain entanglement at higher polymer content, as reported previously^37^.

**Fig. 3).**
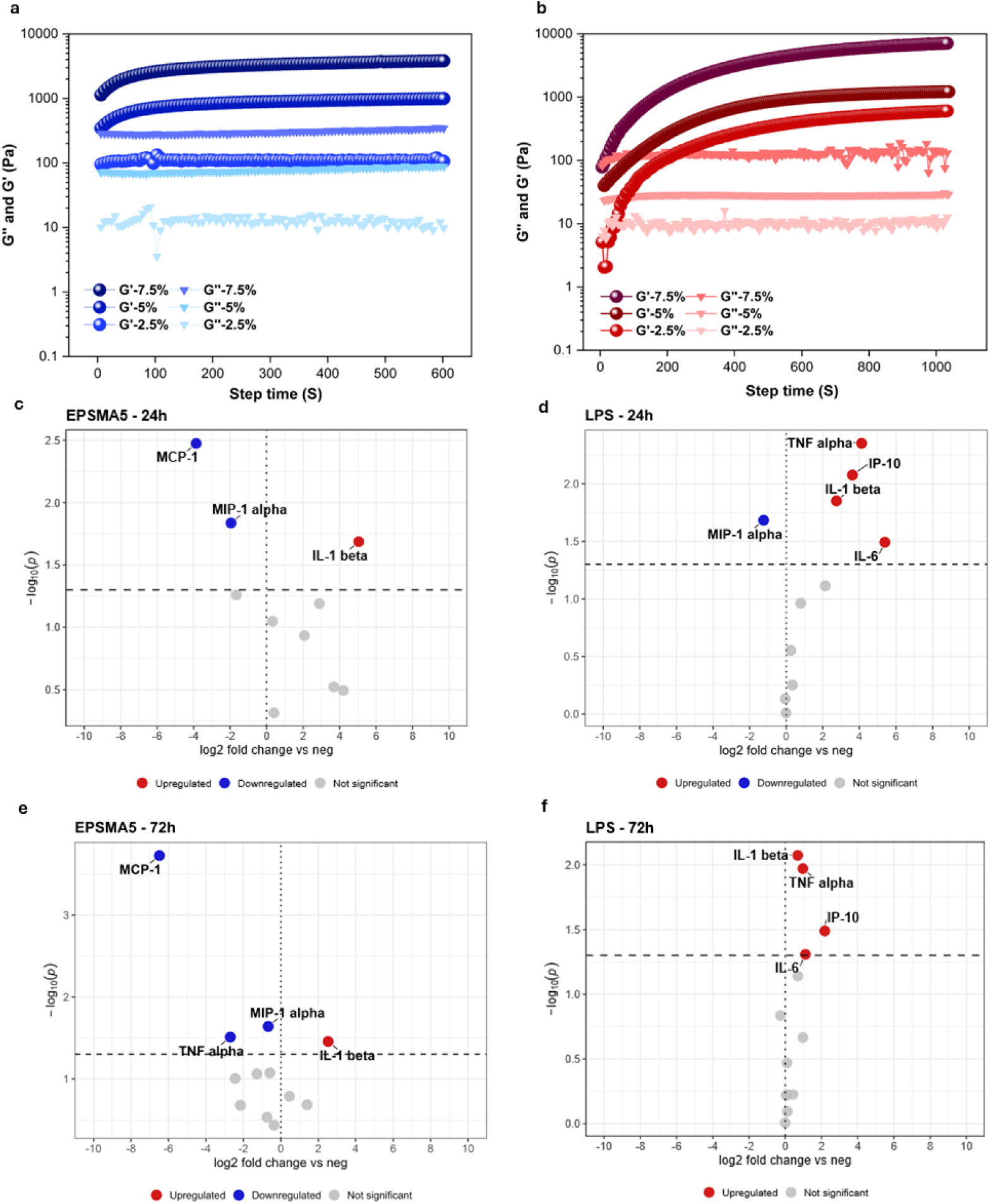
Properties of EPSMA hydrogel. **a**) Rheological properties EPSMA with different concentrations (2.5%, 5%, and 7.5%) under blue light crosslinking (405 nm with 25 mW/cm^2^) **b**) and under red light crosslinking (625 nm with 30 mW/cm^2^). **c,d)** Screening of immunotoxicity effects using THP-1 derived macrophages secreted proinflammatory analytes at 24h and **e,f)** at 72h. Data is shown as volcano plot relative to negative control, unexposed macrophages. P value <0.05 from a Welch’s t-test is considered significant, with analytes above this threshold labelled.

All EPSMA formulations began gelation immediately under blue or red light (G′–G″ crossover). This suggests rapid network formation from pre-formed bacterial biopolymer chains upon irradiation. Blue light showed slightly higher initial G′ and a faster rise due to ambient pre-initiation and rapid radical generation, whereas red light produced slower but more gradual network strengthening, consistent with prior reports^36^.

Additionally, the immunotoxic profile of the EPS was evaluated using a human macrophage proinflammatory cytokine secretion assay. Macrophages were in contact with EPSMA for 24 and 72 h and panels of secreted proinflammatory cytokines from the supernatant were analyzed. Fig. 3c shows that out of all the proinflammatory cytokines tested, EPSMA 5% (w/v) hydrogel only significantly upregulated interleukin (IL) beta secretion. Compared to non-exposed macrophages, EPSMA presence lowers the expression of proinflammatory cytokines MCP-1 and MIP-1 alpha after 24 h. Moreover, this secretion profile remains stable over 72 h of exposure (Fig. 3e). As a comparison, as shown in Fig. 3d,f, we observed the significant increase of several proinflammatory cytokines, namely interleukin-1 beta (IL-1β), tumor necrosis factor alpha (TNF-α), interferon gamma-induced protein 10 (IP-10; CXCL10), and interleukin-6 (IL-6). in lipopolysaccharide (LPS)-treated cells as expected. Unmodified EPS and control biomaterial, nano-fibrillated cellulose, immunotoxicity profiles are shown in supplementary data (Supplementary Note 4 and Fig. S9). A modest, short-term immune response is expected in the case of natural, microbial hydrogels^38^. Moreover, increased stiffness due to crosslinking compared to EPS hydrogel might cause a slight immune response as discussed previously^39^. Additional characterization results, including swelling ratio, microbial contamination assessment, degradability, and shape fidelity have been reported in the Supplementary Material (Note 5 and Fig. S10,11,12).

### Cell viability and 3D-bioprinting

EPSMA cytocompatibility was assessed using 3T3 fibroblasts to confirm its suitability for biofabrication use. Among the tested formulations, 5% EPSMA showed the best performance (>90% viability), with viability slightly higher than the 2D control (Supplementary Note 6 and Fig. S13).

Next, we evaluated the printability of the EPSMA hydrogel using an extrusion-based bioprinter. With optimized parameters, the EPSMA bioink retained its designed geometry after deposition, maintaining structural fidelity in multilayered constructs (Fig. 4a). For comparison, a widely used commercial nanocellulose–alginate bioink was employed as a control (Fig. 4b). Interestingly, EPSMA showed greater swelling and pore-filling behavior compared with the control bioink (Fig. 4c vs 4d) after 6 days of incubation in cell culture medium. In addition, the incorporation of cell-adhesive moieties into the EPSMA hydrogel using an RGD peptide was explored, with the corresponding results reported in the Supplementary Material (Supplementary Note 7 and Fig. S14).

**Fig. 4).**
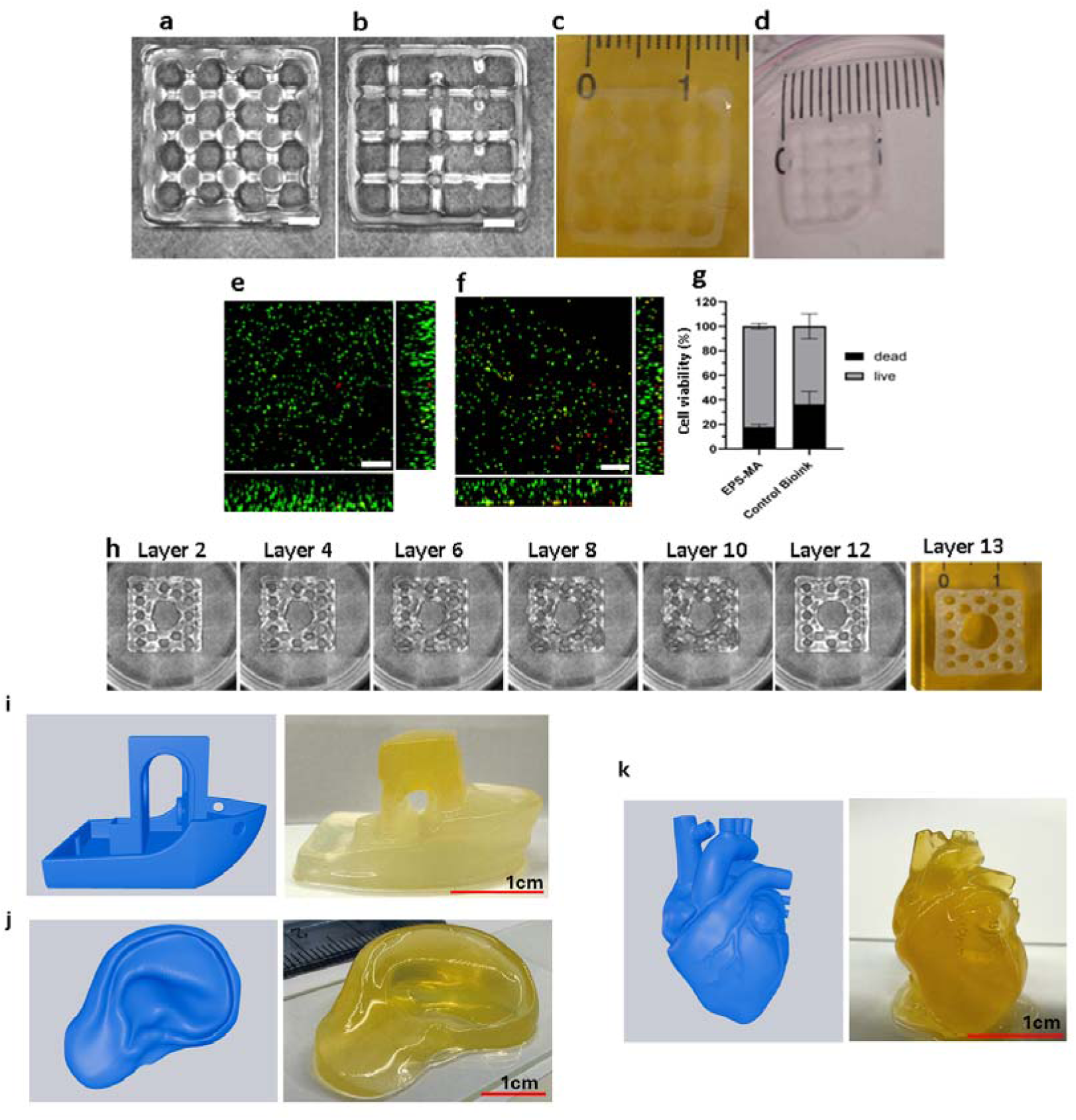
EPSMA as bioink for 3D-Bioprinting. **a**) Cell-laden EPSMA and **b**) Cell-laden control bioink after printing. Totally 6 layers were printed for the structure. **c**) Swelling of EPSMA and **d**) control bioink after 6 days of incubation in cell culture medium. **e**) Representative orthogonal slices of 3T3 cell-laden EPSMA bioink. **f**) and control bioink. Cells were stained with viability/cytotoxicity kit after 6 days of culturing in 3D printed structure. No statistical difference was found between the groups. Scalebar= 200 µm, green for viable cells, red for dead cells. **g**) Percentage of live/dead cells compared to the total number of cells in each sample group. For each biological replicate, at least three regions of interest (ROIs), each containing at least 30 z-sections, were analyzed. **h**) Printability and self-standing properties of 13-layered 1.7 mm thick acellular structure. **i–k**) Proof-of-concept DLP 3D-printing of acellular EPSMA hydrogels (upon 405 nm irradiation and power density of 10 mW/cm^2^): CAD models (blue) and corresponding printed constructs (yellow) for a boat (i), ear (j), and heart (k); curcumin was used as a photoabsorber to improve resolution.

Importantly, when the EPSMA hydrogel was mixed with 3T3 cells and printed into 3D constructs, over 80% cell viability was maintained after 6 days (Fig. 4e, g), comparable to the control bioink (Fig. 4f, g). This level of cytocompatibility is also comparable to widely used bioinks (e.g., alginate and methacrylated gelatin), while the EPSMA offers intrinsic multicomponent complexity that better reflects natural cellular environments. In contrast, alginate is often bioinert and methacrylated gelatin is animal-derived and often needs tuning/blending for robust print fidelity^29,40^. Although EPSMA swelled during culture, the printed structures preserved their overall architecture, underscoring their robustness. To test the ability of EPSMA hydrogels to support complex structures, we further performed acellular extrusion printing to fabricate a multilayer honeycomb design (Fig. 4h). The printed structure retained its resolution across thirteen layers, confirming that EPSMA can form self-standing, thick constructs without collapse. Together, these results support the EPSMA hydrogel as a promising bioink for tissue engineering^41^. Moreover, the EPSMA hydrogel properties can be potentially tuned by modifying cultivation conditions or genetic design to change its composition. As an acellular proof of concept for light-based printing, in addition to extrusion-based 3D-printing, we used a digital light processing (DLP) 3D-printer to construct 3D structures under blue light irradiation (details in the supplementary Fig. S15). Light-based 3D-printing enables the fabrication of hydrogel structures with high geometric complexity and resolution. Three geometries, namely a heart, an ear, and a boat, were printed under blue light (Fig. 4i–k). To improve resolution, curcumin was added as a photoabsorber to attenuate blue light. The resulting acellular constructs showed the gel’s feasibility for light-based 3D-printing. Notably, DLP-printed EPSMA constructs retained their ability to rehydrate and recover their geometry after six months of storage (Fig. S16).

### Injectability and *in situ* crosslinking

Next, we developed a method to monitor hydrogel crosslinking in tissue by validating MB emission in water and 5% EPSMA under red light, which showed a time-dependent signal decrease due to MB reduction (Supplementary Note 8, and Fig. S17a, b).

As a proof of concept for injectability and in situ crosslinking, 5% EPSMA containing MB/TEA/DMA was injected into excised chicken leg muscle (Fig. 5a). Before crosslinking, the low-viscosity precursor was easily injected through a narrow cannula and remained localized at the target site (Fig. 5b, 0 min). After injection, red light illumination was applied (chosen because of its strong soft-tissue penetration and non-invasive initiation), inducing rapid in situ gelation within 20 min at 30 mW/cm² (Fig. 5b). MB emission decreased over time during crosslinking (Supplementary Fig. S18), and dissection confirmed a well-defined hydrogel retained at the injection site (Fig. 5c). Rheological analysis of the recovered gel further confirmed network formation, with the storage modulus remaining higher than the loss modulus over a broad strain range. At high strain, storage modulus decreased and approached loss modulus, indicating gradual network yielding (Fig. 5d). Extrusion results are provided in the supplementary material (Videos S1–S3; Fig. S19).

**Fig. 5).**
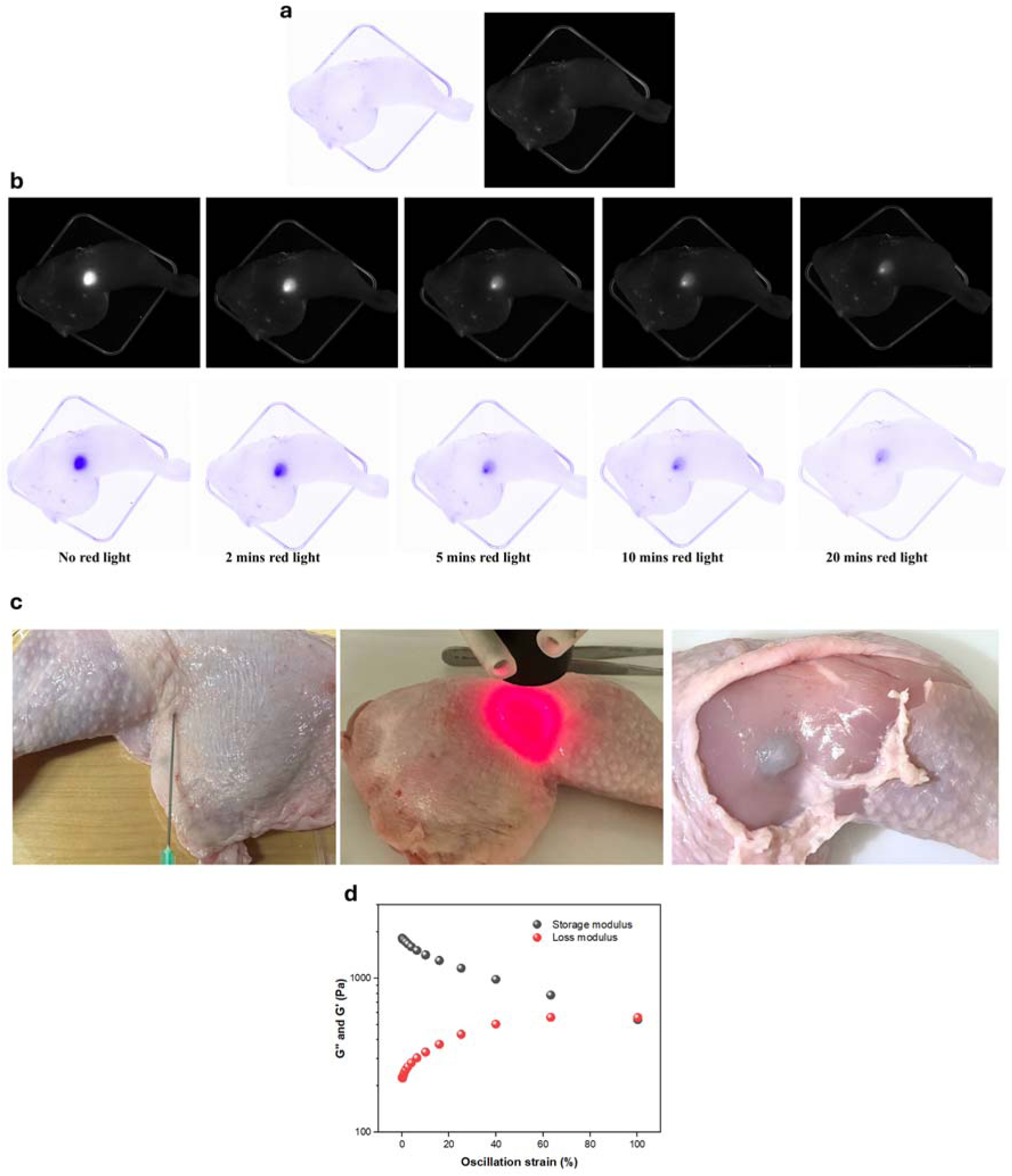
Injectability and in situ red light crosslinking of 5% EPSMA hydrogels monitored by Newton 7 imaging. **a**) Background control: chicken tissue without hydrogel injection. **b**) Tissue immediately after hydrogel injection followed by red light irradiation. **c**) Experimental sequence illustrating hydrogel injection (400 µL) under red light exposure (625 nm with 30 mW/cm^2^ for 20 minutes) and following tissue dissection, revealing the crosslinked hydrogel localized beneath the skin. **d**) Oscillatory strain-sweep rheology of EPSMA after in situ red-light crosslinking within the tissue and subsequent recovery.

This experiment demonstrates that EPSMA can be injected into soft tissue, crosslinked by externally applied red light, and stabilized locally after injection. Thus, it serves as an ex vivo proof of concept for localized and monitorable hydrogel formation through a biological tissue barrier, which may be relevant to minimally invasive drug-delivery and tissue-repair applications. The low-viscosity precursor enables injections, while subsequent crosslinking reduces the risk of displacement before stabilization. In addition, comparison of the gel mass before injection and after tissue dissection showed that approximately 80% of the injected gel could be recovered (Fig. S20), indicating good retention of the material at the target site, with small losses likely occurring during injection, tissue dissection, and gel retrieval. This addresses a practical limitation of injectable photocrosslinkable precursors in dynamic environments, where unpolymerized solutions can otherwise be displaced before curing^42^. Injectable, in situ photocrosslinking hydrogels have been reported previously (including UV/blue light crosslinked methacrylated gelatin)^43^. However, red-light-triggered curing of a bacterially derived hydrogel, combined with MB emission as a readout of crosslinking progression in tissue (Fig. 5b), has rarely been described. This approach enables localized and monitorable hydrogel formation after injection, which may be useful for applications like minimally invasive drug delivery and tissue repair applications.

### Conclusions

This study established EPSMA, a methacrylated EPS produced by ADP1, as a bio-derived, inherently multicomponent hydrogel that combines the molecular richness of bacterial EPS with good cytocompatibility (tested under blue-light crosslinking), injectability, and 3D printability. By optimizing bacterial growth conditions to increase EPS yield and extracting it using only ethanol as the processing solvent, we obtained a multicomponent bacterial hydrogel compatible with both extrusion-based and DLP 3D printing. This intrinsic multicomponent nature provides chemical and functional complexity but may also introduce batch-to-batch variability, particularly in the protein fraction, which should be considered when interpreting and further developing the material. Nevertheless, independent EPS batches used in the 3D-printing and cell viability experiments showed no notable differences in performance. Immunotoxicity assessment indicated only a limited inflammatory response, while residual bacterial components, particularly endotoxins, should be further evaluated. Its dual light-responsiveness further enables *in situ* red light crosslinking after injection, allowing minimally invasive formation and stabilization of 3D constructs within or on tissues. Looking ahead, refining bacterial production and functionalization strategies should broaden the property space and application spectrum of EPSMA hydrogels in biofabrication.

### Experimental methodology

#### Strains and media

*Acinetobacter baylyi* ADP1 (DSM 24193, DSMZ, Germany) was used as the production host for all the experiments. Modified lysogeny broth (LB) agar (LA) plates (10 g/L tryptone, 5 g/L yeast extract, 1 g/L NaCl, and 15 g/L agar) were used to cultivate ADP1 for cell growth and EPS production. D-glucose (10 g/L) was added to the medium unless mentioned otherwise. Kanamycin (30 μg/mL) and 3-azido-2′,3′-dideoxythymidine (AZT; 400 μg/mL) were added to media when required. ABMS medium (Supplementary Note 9) was used for transformation and subsequent cultivation during genetic engineering.

#### EPS production and extraction

For EPS production, overnight precultures were prepared from a -80°C glycerol stock in LB media supplemented with 1% (∼ 50 mM) glucose. The following day, the overnight culture was spread on LA plates supplemented with 1% glucose, unless stated otherwise. The cells were incubated for 72 hours under a range of environmental conditions, including variations in temperature, pH, sugar sources, media compositions, and mineral concentrations. After 72 hours, the cells were harvested from the agar plates and transferred to 50 mL Falcon tubes, then mixed with 16 mL of 0.9% NaCl solution. The mixture was vortexed thoroughly and heated in a water bath at 70-75°C for 15 minutes. The tubes were centrifuged at 13,000 g at 4°C for 15 minutes. The cell-free supernatant was then mixed with 35 mL of 99% cold ethanol and left to incubate in the refrigerator overnight. The following day, the tubes were centrifuged again at 13,000 g at 4°C for 15 minutes, and EPS was collected as a pellet, which was subsequently freeze-dried (ALPHA 1-4 LD-2; Martin Christ Gefriertrocknungsanlagen GmbH, Germany). The dried EPS was weighed immediately after freeze-drying. All samples were processed using three independent biological replicates (n = 3). The results are reported as the mean ± standard deviation (SD), calculated using the STDEV.S function.

#### Genetic engineering

Primers were synthesized by Thermo Scientific (USA), and standard PCR reactions were carried out using Phusion High-Fidelity DNA Polymerase (New England Biolabs, UK). DNA modifications in ADP1 were introduced by natural transformation of recipient strains with linear DNA fragments, as described previously^44^. Briefly, regions of approximately 1000 bp to 2000 bp upstream (left flank (LF)) and downstream (right flank (RF)) of the target genes were amplified from the genomic DNA of ADP1. The LF and RF regions were then assembled together with *tdk-kan^R^* cassette^45^ using splicing by overlap extension (SOE)-PCR and transformed to ADP1, as outlined in previous studies^46^. A rescue cassette was introduced to generate a markerless gene knock-out. Successful gene deletions in ADP1 were selected on LA plates containing 400 μg/mL of AZT and 10 g/L glucose. Genotypes were confirmed by colony PCR using OneTaq (New England Biolabs, UK), and strains were screened for antibiotic sensitivity. Whole-genome sequencing was carried out by Plasmidsaurus (UK). Detailed information on all strains and primers used is provided in supplementary material (Supplementary Table S2 and Table S3).

#### Physical and chemical characterization of EPS

Colorimetric methods were employed to quantify the total protein and polysaccharide content in EPS (UV-Vis spectrophotometer). Briefly, 5-7 mg of dry EPS was dissolved in water. The protein content was determined using the Lowry method^47^, while the polysaccharide content was measured using the phenol–sulfuric acid method^48^. For proteomic analysis, the digested EPS samples were dissolved in 0.1% formic acid, and 800 ng of the sample was injected for analysis. Peptide analysis was carried out using an Easy-nLC1200 system (Thermo Fisher Scientific) coupled to a Q Exactive HF mass spectrometer. Peptides were loaded onto a trapping column and separated on a 15 cm C18 column. The mobile phase consisted of water with 0.1% formic acid (solvent A) and acetonitrile/water (80:20, v/v) with 0.1% formic acid (solvent B). A 90-minute gradient was used, and MS results were collected via Thermo Xcalibur 4.1 software, with data-dependent acquisition (DDA) for MS1 and MS2 scans. The results were analyzed for protein identifications and peptide modifications using MetaMorpheus (version 1.0.6) against the *A*. *baylyi* ADP1 database from NCBI.

To study the monosaccharides in the polysaccharide component of EPS, 5-7 mg of dry EPS was hydrolyzed with trifluoroacetic acid in a 100°C water bath for 6 hours. After hydrolysis, the sample was dried using methanol and nitrogen gas. The remaining residues were dissolved in ultrapure water (Milli-Q system) and filtered. The filtered sample was then analyzed for its monosaccharide composition using an LC-20 AC Prominence liquid chromatograph (Shimadzu, USA) equipped with a RID-10A refractive index detector. Separation was carried out on a Phenomenex Rezex RHM-monosaccharide H (8%) column, with ultrapure water as the mobile phase at a flow rate of 0.5 mL/min and a column temperature of 40°C.

DNA was extracted from EPS using a phenol:chloroform:isoamyl alcohol protocol, precipitated with sodium acetate and isopropanol, washed with ethanol, and resuspended in Tris–EDTA buffer as previously described^49^. DNA concentration was determined by Qubit fluorometric quantification. Fourier transform infrared spectroscopy (FTIR; Spectrum Two, PerkinElmer Inc., Waltham, MA, USA) was applied to identify the functional groups present in EPS, allowing for a better understanding of its chemical composition.

Field-emission scanning electron microscope (FESEM; UltraPlus, Carl Zeiss Microscopy GmbH, Germany) was used to investigate the surface morphology of EPS. Additionally, a rheometer (Discovery HR-2, TA Instrument, USA) was employed to assess the mechanical properties of EPS by measuring the storage modulus, loss modulus, viscosity, and adhesiveness of EPS. A tack test was performed using a Discovery HR-2, TA Instrument to evaluate tissue adhesiveness of EPS-hydrogels^50^. Two freshly excised chicken breast tissues were mounted on the upper and lower plates of the rheometer. A small volume (∼0.3–0.5 mL) of hydrogel was dispensed between the tissues and compressed at 0.1 N for 60 s to ensure wetting. The test was performed in tensile (normal force) mode, where the upper plate was then retracted axially at 20 μm s ¹ while the normal force was continuously recorded. Two hydrogel formulations (one glucose-based (Glu-Gel) and one rhamnose-based (R-Gel)) were tested under identical conditions at room temperature. The separation behavior was recorded to assess both adhesive (interface) and cohesive (gel-body) failure modes.

#### Methacrylation and crosslinking of EPS with blue and red light

Details of the materials used for methacrylation and photocrosslinking are provided in the supplementary material (Supplementary Note 10). Methacrylation of EPS-hydrogel was carried out through a nucleophilic substitution reaction, where the free amine and hydroxyl groups of the polypeptide reacted with methacrylic anhydride. First, 600 mg of EPS-hydrogel was dissolved in 170 mL of deionized (DI) water at room temperature (RT) under constant stirring until a clear solution was obtained. Once fully dissolved, 900 μL of methacrylic anhydride was added to the solution in three equal portions. After each addition, the reaction mixture was stirred for 30 minutes at RT to allow methacryloylation to proceed. The pH of the solution was carefully readjusted to 7.5–8.0 using 1 N NaOH after each addition, since the reaction tends to lower the pH. Following the final addition, the reaction mixture was stirred continuously at RT for 6 hours. During this period, the pH was monitored and adjusted every 15 minutes to maintain it between 7.5–8.0. After 6 hours, the pH stabilized and then it was adjusted one final time to 7.5, and the mixture was left to stir overnight at RT to ensure complete reaction. The next day, the reaction solution was transferred to a dialysis membrane (MWCO 3.5 kDa) and dialyzed against DI water at RT for 4 days to remove unreacted reagents and byproducts. The dialysis water was replaced twice daily to maintain efficiency. After purification, the methacrylated product was recovered by lyophilization and stored at 4 °C until further use. The methacrylated EPS gel was named EPSMA in this study. The degree of methacrylation was quantified by the trinitrobenzene sulfonic acid (TNBS) assay and analyzed using a UV−vis spectrophotometer (Shimadzu UV-3600 Plus). To confirm the methacrylation qualitatively, ¹H-NMR spectroscopy was performed in D O using a 500 MHz NMR spectrometer (SCZ500R, JEOL Resonance, Japan).

After methacrylation, for blue light crosslinking, Lithium phenyl-2,4,6- trimethylbenzoylphosphinate (LAP) was used at the final concentration of 0.3%. For red light crosslinking, Methylene blue (MB+), Triethanolamine (TEA), and N,N-Dimethylacrylamide (DMA, 99%, contains 500 ppm monomethyl ether hydroquinone as inhibitor) were used at final concentrations of 1.6%, 1%, and 4%, respectively. The inhibitor was then removed by passing through a column of aluminum oxide. For red light in situ photorheology, a Thorlabs LED (625 nm, M625L4, 30 mW/cm^2^ and for blue light *in situ* photorheology, a Thorlabs LED (405 nm, M4405L4, 25 mW/cm^2^) equipped with a collimator head was placed at 7 cm from the parallel plate for irradiating the sample during the measurement. All rheology measurements were performed using a rotational rheometer (Discovery HR-2, TA Instruments Inc., USA) in a parallel plate geometry with a diameter of 12 mm and the measurement gap was set to 2.1 mm. For rheological measurements, a total sample volume of 300 µL was placed on the rheometer plate.

The swelling behavior of 5% (w/v) EPSMA hydrogels containing 0.3% (w/v) LAP was evaluated in 3T3 growth medium at 37 °C. After photocrosslinking, the initial mass of each hydrogel was recorded. At predetermined time points, the samples were removed from the medium, gently blotted to remove excess surface liquid, weighed, and returned to the medium. The swelling ratio was calculated as the percentage increase in hydrogel mass relative to its initial mass. For microbial contamination assessment, EPSMA samples were spread onto LA plates supplemented with glucose and monitored for colony formation. Sterile water and ADP1 cells were used as negative and positive controls, respectively. For shape fidelity and filament diameter images of independently extrusion-based 3D-printed constructs were analyzed using ImageJ. The image scale was calibrated using a known dimension. Filament diameter was measured perpendicular to the filament direction at multiple positions. Shape fidelity was evaluated by comparing the measured dimensions of the printed structures with the corresponding designed dimensions.

For enzymatic degradation test, three independently prepared 5% (w/v) EPSMA hydrogels containing 0.3% (w/v) LAP were photocrosslinked using 405 nm light for 10 min. The hydrogels were then incubated in PBS at 37 °C either without enzyme or in the presence of 0.1 mg/mL Proteinase K or 1 mg/mL lysozyme. The wet weight of each sample was measured daily after gently removing excess surface liquid. After each measurement, the samples were returned to fresh medium. Changes in hydrogel weight were expressed relative to the initial wet weight. Three independently printed constructs were analyzed, and the results are reported as mean ± SD.

#### Immunotoxicity assay

Human THP-1 monocytes (ATCC TIB-202, American Type Culture Collection, USA) were maintained in RPMI-1640 medium (Gibco) supplemented with 10% heat-inactivated fetal bovine serum (FBS; Gibco). Cells were cultured under standard conditions at 37 °C in a humidified atmosphere containing 5% CO and passaged according to established laboratory procedures. THP-1 cells were subcultured for at least two passages following thawing before use in experiments. To induce macrophage-like differentiation, THP-1 cells were cultured in RPMI-1640 supplemented with 10% heat-inactivated FBS and 25 nM phorbol 12-myristate 13-acetate (PMA; Sigma-Aldrich). Cells were seeded into T75 flasks at a density of approximately 1.25 × 10 cells cm ² and incubated for 48 ± 2 h. Following differentiation, the activation medium was removed, and cells were detached using TrypLE Express (Gibco). Differentiated THP-1 macrophage-like cells were seeded into 96-well plates at a density of approximately 4.0 × 10 cells per well in 50 μL culture medium. Biomaterials were prepared directly in the wells prior to cell seeding. Methacrylated EPS (EPSMA 5%) hydrogel was photocrosslinked (1 minute under UV light) prior to cell exposure. In addition, nanofibrillar cellulose (NFC, 1%, GrowDex-B, UMP Kymmene Oyj), known to be non-inflammatory gel, was used as 3D negative control. Lipopolysaccharide (LPS, 100 ng mL ¹; Sigma-Aldrich) was included as a positive pro-inflammatory control. Cells were exposed to the test materials for either 24 or 72 h. Three technical replicate wells were included for each experimental condition. Following exposure, culture supernatants were collected and analyzed for cytokine secretion using a multiplex immunoassay (ProcartaPlex custom-made assay, Thermo Fisher Scientific) according to the manufacturer’s instructions. Selected pro-inflammatory cytokines: CXCL11, IFN-gamma, IL-1 alpha, IL-1 beta, IL-17A, IL-18, IL-6, IL-8, IP-10, MCP-1, MIG, MIP-1 alpha, and TNF alpha were analyzed from cell culture supernatant. Results were visualized and analyzed using in-house built R-script as described previously^51^. Data are presented as mean ± standard deviation (SD). Comparisons between groups are shown as log2 fold change compared to negative control (differentiated THP-1 cells without exposure) and statistical analysis was performed using a Welch’s *t*-test. Differences were considered statistically significant at *p* < 0.05.

#### Cell viability

The viability of the 3T3-Swiss albino fibroblastic cells (3T3, ATCC, CCL-92) inside EPSMA scaffolds was evaluated. The cells were cultured in 3T3 growth medium (3T3GM) containing Dulbecco’s Minimum Essential Medium (DMDM, Gibco, 21969035), Foetal bovine serum (FBS, Sigma-Aldrich, F7524), GlutaMAX Supplement (Gibco, 35050061), and Penicillin/streptomycin (PS, 15140122). Cells were cryopreserved in 3T3GM+10% dimethyl sulfoxide (DMSO, Sigma, D5879) in 1*10^6 cells/vial. Cells were thawed into T75 flasks and passaged 3 times before using in experiments. 3T3GM was changed 3 times per week and cell detachment was done using Tryple Select (Gibco, 12563-029). For the experiments, detached cells were counted and 3 milllion cells/mL were used to encapsulate inside EPSMA hydrogels. For the viability tests, cell-laden EPSMA-hydrogel was pipetted into 96 well plate, ∼50µL per well. 7.5%, 5%, and 2.5% EPSMA material were used for the viability experiment. EPSMA gels were exposed to 365 nm UV light for 2 mins. 2D control cells were plated on top of cell culture treated polystyrene well plate on density of 3*10^4 cells/cm^2^. To evaluate the effect of UV-exposure on cell viability, half of the control cells were exposed to same UV light (365 nm) exposure as crosslinked gels. The viability of the cells inside EPSMA-based hydrogels was evaluated after 6 days inside the hydrogel. Cell viability was observed using live/dead viability/cytotoxicity kit (Invitrogen, L3224) according to the manufacturer’s instructions. Briefly, 0.5µM Calcein AM + 0.25µM Ethidium homodimer working solution was prepared in Phosphate Buffered Saline (PBS). The solution was pipetted into the wells, 100 µL per well and incubated for 30 minutes. Thereafter, the wells were imaged using Zeiss LSM 800 confocal microscope (Carl Zeiss Microscopy GmbH, Germany). To further investigate cell adhesion within EPSMA hydrogels, an RGD peptide was physically mixed with EPSMA at a final concentration of 0.1 mg/mL prior to hydrogel crosslinking. 3T3 fibroblasts were encapsulated in either RGD-containing EPSMA (RGD-EPSMA) or EPSMA without RGD and cultured for 6 days. Cells cultured in 2D without any hydrogel were included as a control. Cell viability was then evaluated using the Live/Dead viability/cytotoxicity assay following the same staining and imaging procedure described above. All samples were processed using three independent biological replicates (n = 3). The results are reported as the mean ± standard deviation (SD), calculated using the STDEV.S function.

#### 3D-printing of EPSMA without cells

EPSMA hydrogel was diluted to 5% (w/v) and mixed with 0.3% (w/v) LAP. PBS was used to adjust the formulation to the required concentrations. The ink was printed using a 3D Bioplotter (EnvisionTEC GmbH, Gladbeck, Germany) equipped with a low-temperature printhead and a 25 G blunt-end needle (CELLINK, NZ6300505001), at a pressure of 1.2 bar and a printing speed of 4.0 mm/s. A 10 × 10 mm six-layer grid structure and a 13-layer honeycomb structure were printed to evaluate printability and post-printing structural stability. Each deposited layer was photocrosslinked for 5 s, followed by a final 60 s exposure of the complete printed structure. Following printing, the structures were incubated at 37 °C in a humidified atmosphere containing 5% CO .

Digital light processing (DLP)-based 3D printing was performed using a Creality LD-002R printer equipped with a custom-made vat (Supplementary Fig. S10). The exposure time was 30 s per layer using 405 nm light at an intensity of 10 mW/cm², and the layer thickness was 50 µm. The printing formulation consisted of 2% (w/v) EPSMA, 0.5% (w/v) LAP, 4% (v/v) DMA, and 1% (w/v) curcumin purchased from Sigma-Aldrich.

#### 3D-bioprinting of cell-laden EPSMA bioink

For cell-laden bioinks, 3T3 cells were mixed thoroughly with 5% (w/v) EPSMA at a cell density of 5 × 10 cells/mL. The prepared cell-laden EPSMA bioink was transferred into printing cartridges and extruded through a 25 G blunt-end needle at a pressure of 1.1 bar and a printing speed of 5.0 mm/s. Each deposited layer was photocrosslinked for 5 s, followed by a final 60 s exposure of the complete printed structure. As a control, 3T3 cells were incorporated into CELLINK Bioink at the same cell density and printed under the corresponding conditions.

Cell viability was evaluated using a LIVE/DEAD Viability/Cytotoxicity Kit (Invitrogen, L3224) according to the manufacturer’s instructions. The samples were imaged using a Zeiss LSM 800 confocal microscope. For each sample, at least three regions of interest containing at least 30 z-sections were analyzed. The proportion of dead cells relative to the total cell count was determined using an in-house ImageJ macro (National Institutes of Health, version 1.54), and the data were plotted using Prism (GraphPad, version 10.3.1). Statistical significance between groups was evaluated using one-way ANOVA, with p < 0.05 considered statistically significant. All samples were processed using three independent biological replicates (n = 3). The results are reported as the mean ± standard deviation (SD), calculated using the STDEV.S function.

#### Injectability and *in situ* crosslinking of EPSMA-hydrogel

For injectability and *in situ* crosslinking analysis, EPSMA-hydrogels were prepared at 5% (w/v) concentration. The MB (1.6%), DMA (4%), and TEA (1%) were added to the hydrogel for red light crosslinking. Control samples included pure water and water containing MB, DMA, and TEA to assess background emissions. The emission signal of MB was recorded using a *Vilber Newton 7* imaging instrument. Chicken leg muscle tissue was obtained fresh from a local food source and used within the same day. The hydrogel precursor was loaded into a 1 mL syringe fitted with a 21G cannula. Approximately 400 µL of precursor solution was injected under skin. Following injection, the tissue was exposed to red light (λ ≈ 625 nm) for 20 minutes. Images were taken at 0, 2, 5, 10, and 20 minutes post-irradiation to document the changes in hydrogel appearance and emission. After irradiation, tissue samples were dissected to visualize the crosslinked hydrogel at the injection site. Additionally, injectability was assessed using Instron 4411 mechanical testing device (Instron Corp., USA) equipped with 500 N load cell and custom compression head was combined in compression mode with sterile 1 mL plastic luer-lock syringe (Fisher Scientific, USA) and 25G needle (BD Microlance™ 3, Becton, Dickinson and Company, New Jersey,USA). A 1 mL syringe filled with the 5% EPSMA-hydrogel and the mechanical testing device pressed the syringe plunger 16 mm at three speeds (200, 400, and 500 mm/s), and the required force (N) against the displacement (mm) was recorded. Data was acquired with Instron Series IX software (Instron Corp., USA).

#### Computational details

All atom molecular dynamics simulations were used to study the structural and dynamical effects of glycan moieties on the protein, both in solution and in the covalently glycosylated form. Protein structures were generated using AlphaFold and CHARMM-GUI, parameterized with the ff14SB and GLYCAM06 force fields, and simulated using AMBER22. Standard force fields and simulation protocols were applied, followed by long production runs to capture protein-glycan interactions. Full methodological details are provided in the Supporting Information section (Supplementary Note 1).

## Data availability

The data of this study are available from the corresponding author upon reasonable request.

## Author contributions

S.R.: conceptualization, methodology, investigation, formal analysis, writing. P.C.I., N.P.: Investigation. A.E.: methodology, investigation, formal analysis, writing. L.Y.-O.: conceptualization, methodology, investigation, formal analysis, writing. V.S.P., M.Kh., J.M.: methodology, investigation, formal analysis. D.G., T.L., M.K., H.S.: supervision, resources, funding acquisition. S.S., V.S.: conceptualization, supervision, resources, funding acquisition. All authors: review and editing of the manuscript.

## Acknowledgment

SR would like to thank the Suomen Kulttuurirahasto Foundation (05232023). SS would like to thank the Novo Nordisk Foundation (grant NNF21OC0067758) and the Research Council of Finland (grant no. 347204, 353587, and 372132). VS would like to thank the Novo Nordisk Foundation (grant NNF21OC0079579) and Research Council of Finland (367615). This research was also supported from the European Union – NextGenerationEU instrument and is funded by the Research Council of Finland under grant number 353658. MK and VP would like to thank the Research Council of Finland project 353173 (Center of Excellence Body-on-Chip) and SUSBIO funding (352754). TL and AE would like to thank the European Research Council (ERC Consolidator Grant Project PADRE, No 101001016) and Tampere Institute for Advanced Study. DG would like to thank the European Research Council (ERC Consolidator Grant ARCHIMEDES, No. 101043848).

## Competing Interests

A patent application related to this work has been filed.

## Notes

### Summary of Updates

This version corrects a figure assembly error in the previous version. The manuscript has been updated to include the correct Figure 5.

