## Supplementary material for "3D-Printable and Cytocompatible Hydrogel from *Acinetobacter baylyi* ADP1 Extracellular Matrix"

Table S1. The 10 most abundant proteins in EPS matrix based on proteomic analysis.

| **Protein Descriptions** | **Predicted location** |
|---|---|
| putative biofilm related protein | Surface/extracellular |
| conserved hypothetical protein/Tetratricopeptide repeat family protein | Periplasm /extracellular |
| putative Outer membrane protein precursor (OmpA-like) | Outer membrane |
| putative hemagglutinin/hemolysin-related protein hemagglutinin repeat-containing protein | Extracellular |
| conserved hypothetical protein\| putative membrane protein | Inner/outer membrane |
| putative phosphate transporter (ABC superfamily, peri-bind) | Periplasm |
| outer membrane protein (AdeC-like) | Outer membrane |
| putative outer membrane protein | Outer membrane |
| Protein AcuA\| putative fimbrial-like protein | Surface/ outer membrane |

### Note 1: Computational details

All-atom molecular dynamics (MD) simulations were carried out to investigate the behavior of the protein in the presence of glycan moieties. The initial protein structure was generated using AlphaFold[1], and its conformational dynamics were examined both in the absence and presence of HexNAc at concentrations ranging from 0 to 0.1 M. In addition to these solution-phase simulations, the protein was covalently functionalized with the corresponding glycan groups, and the structural and dynamical properties of the resulting glycoprotein were analyzed. The glycoprotein model was constructed using CHARMM-GUI[2].

Protein atoms were described using the ff14SB force field[3], while the glycan groups were parameterized with the GLYCAM06 force field[4]. The protein systems, including both the free protein and the glycosylated complexes, were centered in a rectangular periodic simulation box with dimensions of 19 × 19 × 27 $nm^3$. Periodic boundary conditions were applied in all directions. Each system was solvated using the TIP3P water model[5], and appropriate numbers of $Na^+$ and $Cl^-$ ions were added to neutralize the total system charge.

All simulations were performed using AMBER22[6]. Prior to dynamics, each system was subjected to 100,000 steps of energy minimization to eliminate steric clashes and unfavorable contacts. The minimized systems were then slowly heated from 0 K to 300 K under constant-volume (NVT) conditions. Temperature control was achieved using a Langevin thermostat with a collision frequency of 1 $ps^{-1}$[7]. During the heating stage, harmonic positional restraints with a force constant of 2.5 $kcal \cdot mol^{-1} \cdot Å^{-2}$ were applied to the protein and ligand atoms. This phase was carried out for 2 ns using a 1 fs integration time step. Following heating, the systems were equilibrated for an additional 5 ns under constant-pressure (NPT) conditions at 300 K and 1 bar, maintaining a 1 fs time step. Pressure coupling was handled using an isotropic Berendsen barostat with a relaxation time of 1 ps[8]. Subsequently, all positional restraints were removed, and production MD simulations were performed for 150 ns with a 2 fs time step to characterize the interactions between the glycan groups and the protein. Long-range electrostatic interactions were treated using the Particle Mesh Ewald (PME) method with a real-space cutoff of 12 Å[9]. All covalent bonds involving hydrogen atoms were constrained using the SHAKE algorithm[10].

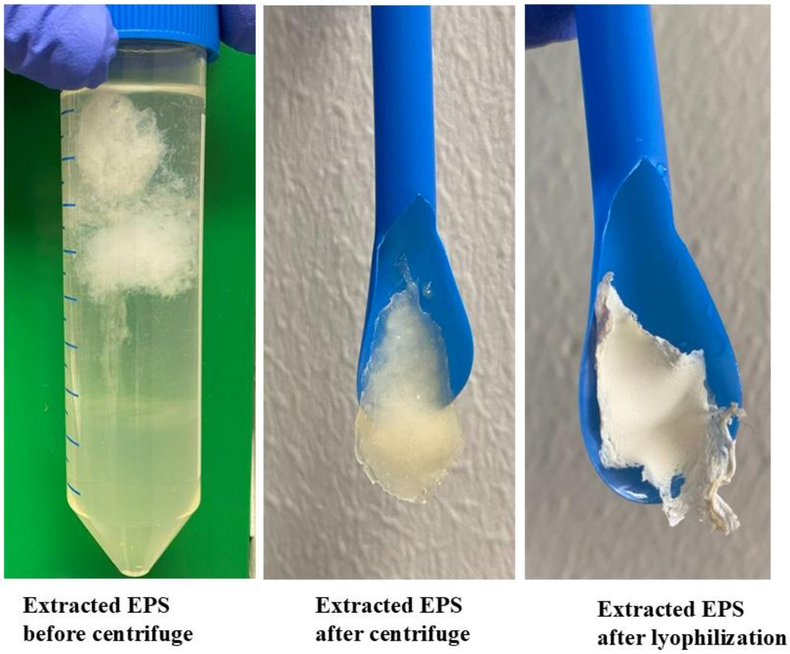


**Fig. S1.** Recovery of ADP1-derived EPS during extraction. Representative images showing the EPS-rich suspension before centrifuge, the gel-like EPS recovered after centrifuge, and the final freeze-dried EPS material used for further hydrogel preparation.

### Note 2: Role of c-di-GMP producer genes in EPS production

All seven c-di-GMP genes could be deleted individually, but during construction of the strain with all 7 genes deletions, gene ACIAD2242 consistently resisted removal, and we ended up with ASA1416. Whole-genome sequencing confirmed successful deletion of the other six genes. Under the same growth conditions, deletion of each gene individually, as well as combined removal of all six, did not result in significant changes in EPS yield compared with wild type. (Fig. S1a). While small fluctuations were observed across mutants, they were within the range of biological variability. These results contrast with systems such as cellulose synthase and PNAG synthase, which are activated by c-di-GMP[11,12]. Instead, our results indicate that ADP1 EPS production is largely insulated from global changes in diguanylate cyclase content. One possible explanation could be related to EPS biosynthesis/export machinery. The ADP1 EPS is polymerized by a Wzx/Wzy-type pathway and exported through a Wza-like pore. Regulation of such systems is thought to occur through tyrosine kinase–phosphatase modules[13]. Because no PilZ-type or c-di-GMP–binding effector is known in this pathway, it is plausible that Wza-dependent polymers are not directly sensitive to c-di-GMP. Additionally, any single knockout, or even combinations of knockouts, did not result in significant growth defects, suggesting that they are not required for core growth (Fig. S2a, b).

To explore whether GGDEF proteins might act only under environmental stress, we compared EPS production across two physiologically relevant parameters: temperature and pH. Wild-type cells showed condition-dependent changes (Fig. S3). However, none of the GGDEF knockouts displayed major pattern differences from wild type across these environments. We also examined EPS variation within each knockout strain across conditions to test whether loss of a given GGDEF would flatten the response profile. CHASE8-domain protein (ASA1400) and the PAS-domain protein (ASA1402) deletion showed reduced EPS variation between pH values, suggesting they may play roles in pH-specific regulation. Overall, these results suggest that in ADP1, GGDEF enzymes do not exert global control over EPS yield, even under environmental conditions that modulate wild-type production. Instead, their modular sensory domains and hybrid GGDEF–EAL architectures are consistent with local signaling roles, producing c-di-GMP microdomains that may regulate specific extracellular proteins, adhesins, or pili, rather than the polysaccharide backbone itself. Given that ~75% of the EPS mass is polysaccharide and only ~20% is protein, even substantial changes in protein-linked regulation would not be expected to significantly alter net EPS yield. Instead, c-di-GMP may shape matrix architecture, adhesiveness, and mechanics—phenotypes not captured by bulk EPS quantification.

Notably, the ASA1416 strain showed pronounced aggregation even at 18 h compared to wild type (Fig. S4). This may reflect that c-di-GMP influences cell motility and surface separation, such that loss of multiple GGDEFs promotes early flocculation. Alternatively, c-di-GMP signaling may alter EPS composition or distribution, shifting it toward a more capsular and adhesive state rather than freely released polymer, thereby enhancing cell–cell adhesion and aggregation despite unchanged bulk EPS yield.

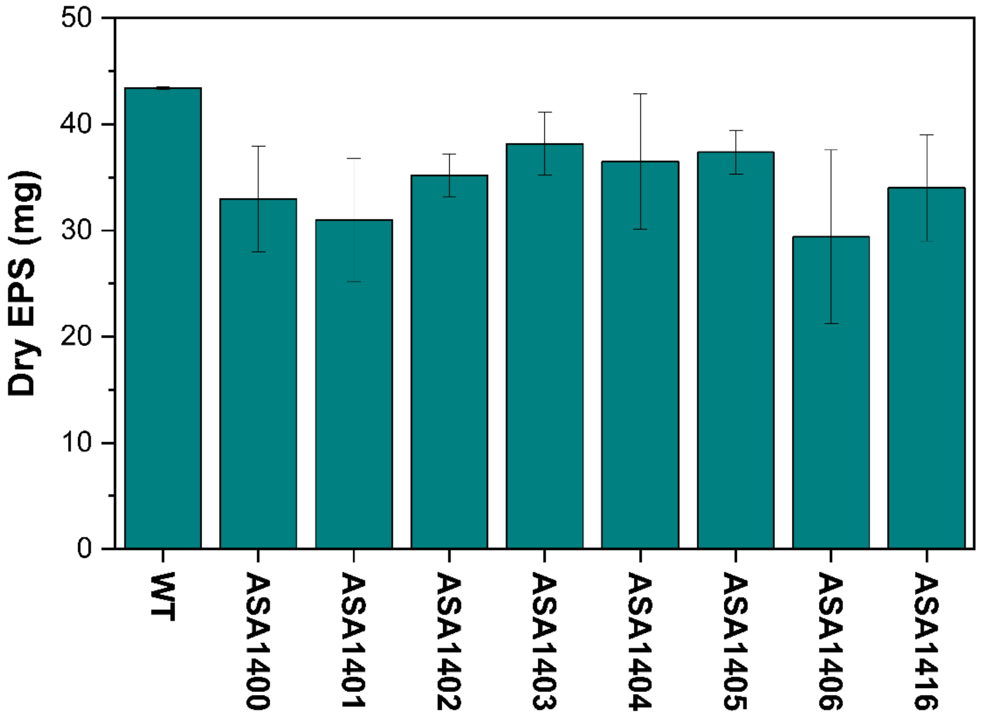


Fig. S2. EPS production yield by WT-ADP1 compared with deleted c-di-GMP producing genes.

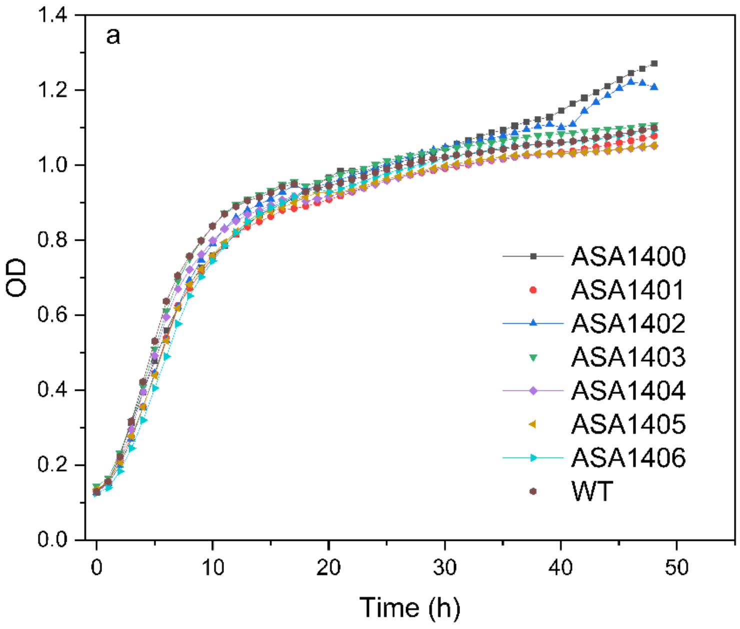


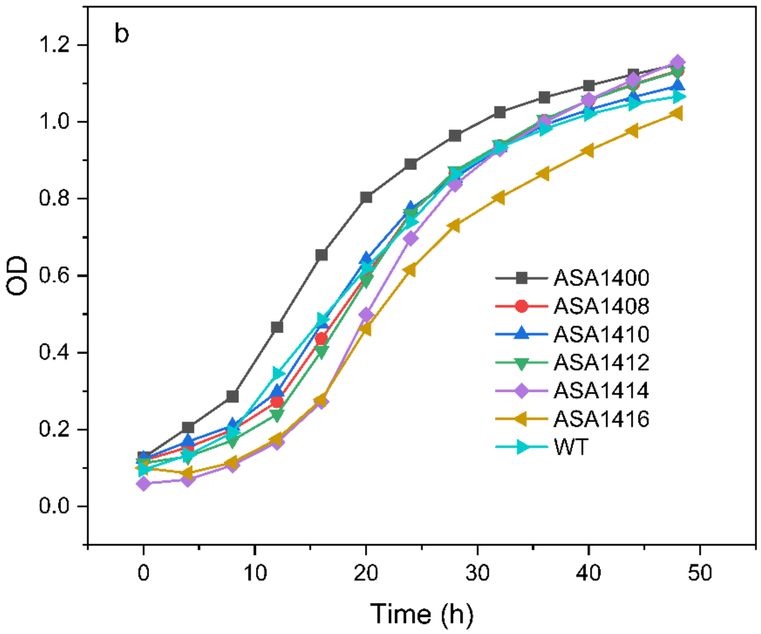


Fig. S3. Optical density (OD) of cultures in LB-Glu1% media. a) Growth rate of single gene deleted strains compared with ADP1-WT. b) Growth rate of multipele genes deleted strains compared with ADP1-WT. Data are presented based on three biological replicates.

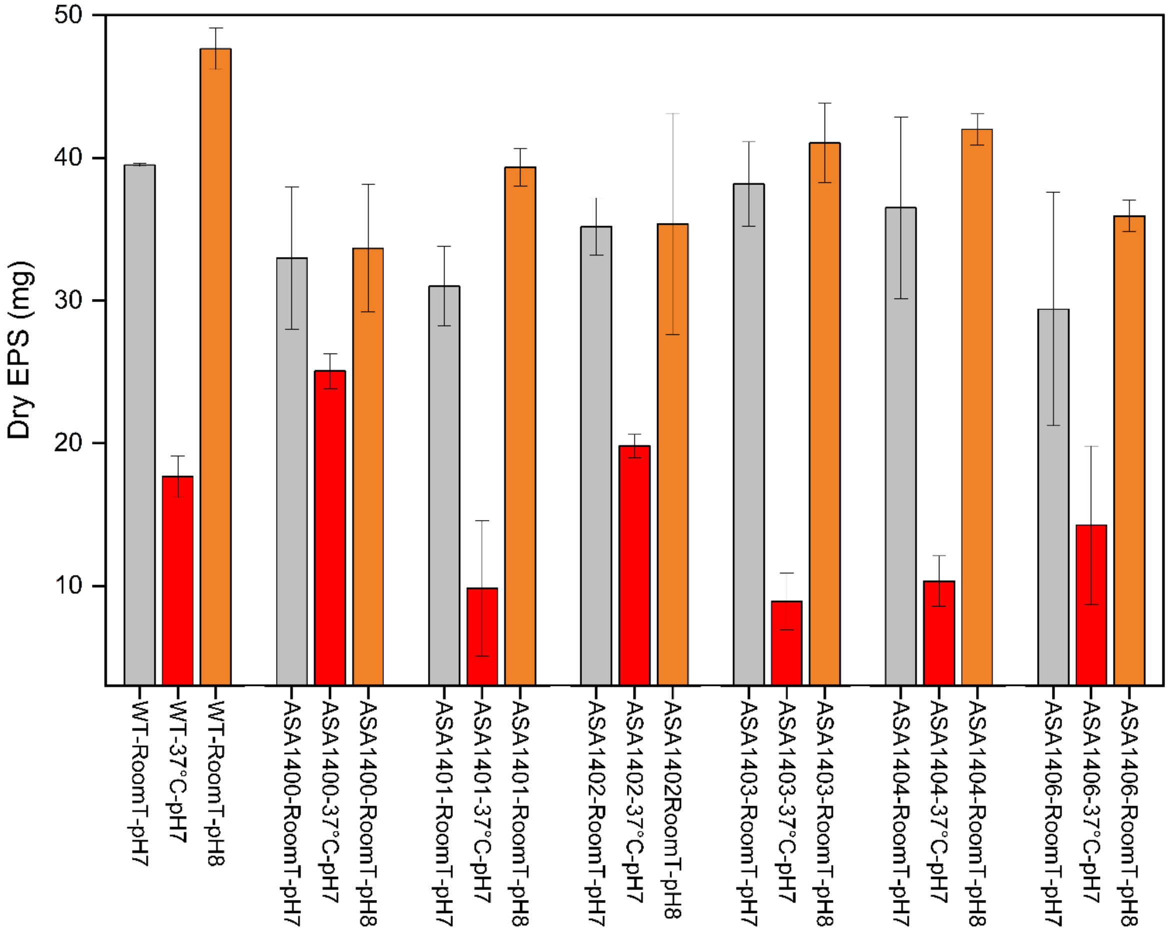


Fig. S4. EPS yield of single-gene deletion strains across different environmental conditions. Data are presented based on three biological replicates.

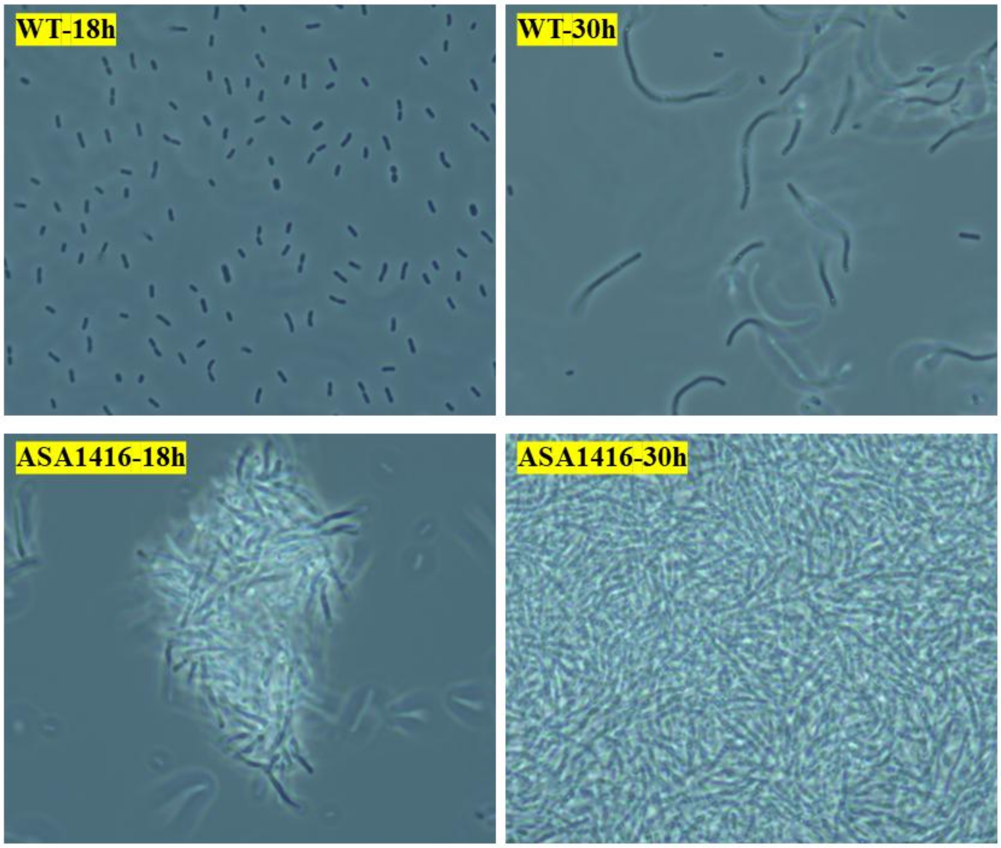


Fig. S5. Microscopic images of WT-ADP1 compared with ASA1416 strain.

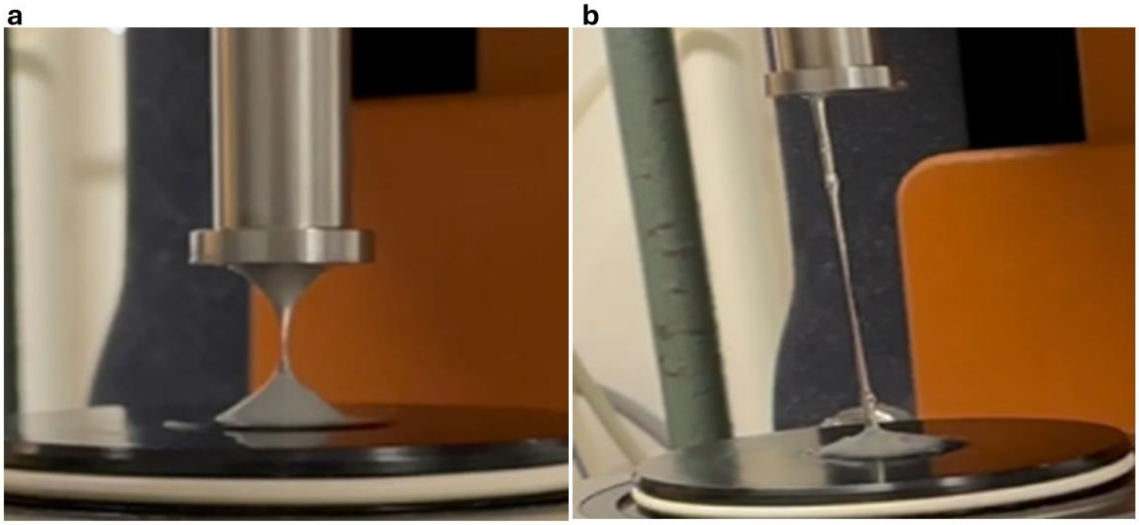


Fig. S6. The hydrogel adhesive behavior during probe-tack test on a rheometer. a) Glucose-based EPS hydrogel. b) Rhamnose-based EPS hydrogel.

**Note 3: Methacrylation of EPS hydrogel**

The degree of methacrylation of EPSMA (methacrylated EPS), based on amine modification, was determined to be ~45 ± 5%, with minor batch-to-batch variations This value was quantified using the TNBSA assay, based on UV–vis absorption measurements. The calculation of methacrylation degree was derived from the quantification of free amine groups in the EPSMA hydrogel relative to the unmodified EPS hydrogel. It is important to note that this method accounts only for modifications at the amine functionalities. Since the EPS hydrogel contains polysaccharide components, methacrylation of hydroxyl groups on the EPS backbone is not captured by the TNBS assay, and therefore the reported values represent a partial estimation of the overall modification. To further validate the chemical modification, $^1$H-NMR spectroscopy was performed. The spectra of the EPSMA showed distinct olefinic proton peaks at 5.5 and 6.15 ppm corresponding to methacryloyl groups (Fig. S6), which were absent in the spectra of the unmodified EPS (Fig. S7). The appearance of these additional peaks qualitatively confirmed the successful incorporation of methacryloyl groups into the EPS structure.

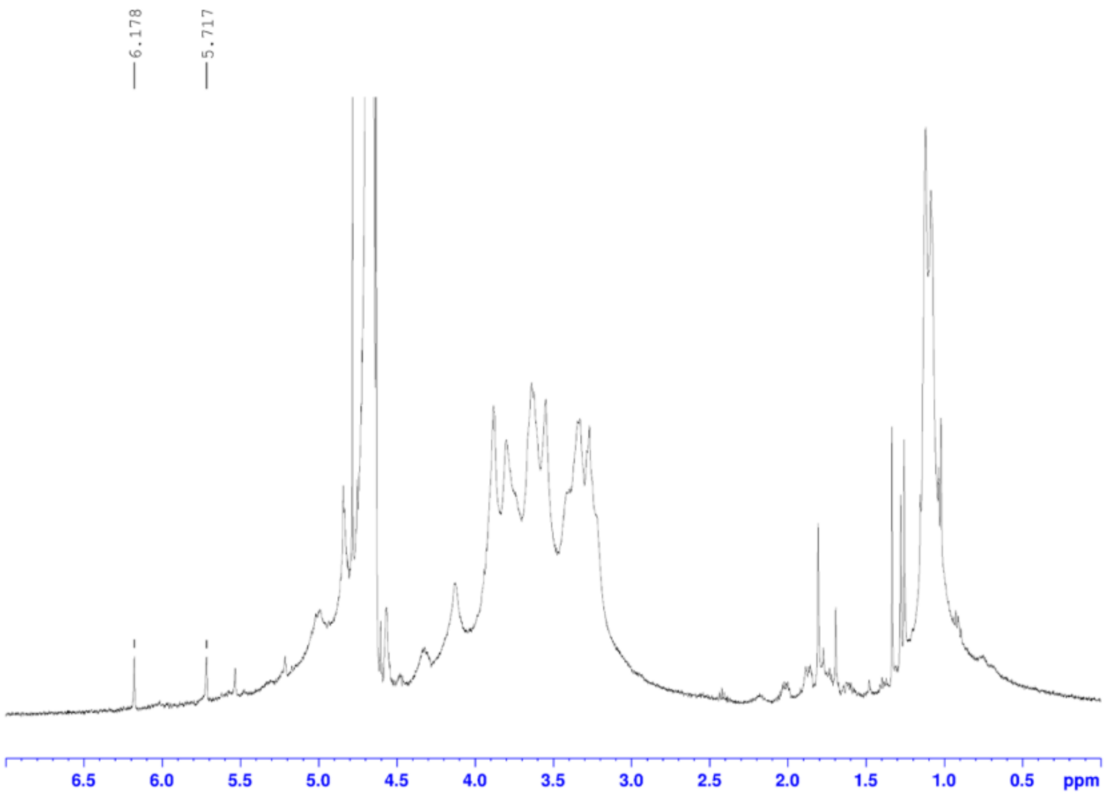


Fig. S7. $^1$H-NMR (500 MHz) spectrum of methacrylated EPS (EPSMA) in $D_2O$.

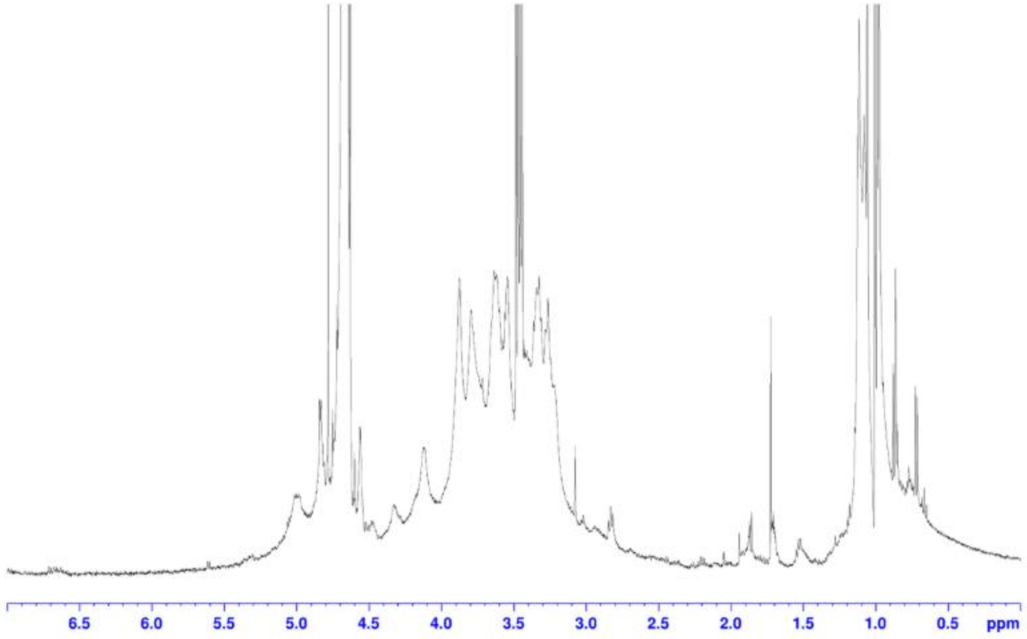


**Fig. S8.** $^{1}$H-NMR (500 MHz) spectrum of pure EPS in $D_2O$.

**Note 4:** At 24 h, EPS5 (pure EPS 5%) induced significant increases in TNF-α and IL-1β, while MCP-1 was reduced compared to negative control (unexposed activated macrophages). After 72 h, however, the response shifted predominantly toward reduced secretion of several pro-inflammatory analytes, including MCP-1, IFN-γ, IL-1α, IL-8, MIP-1α, and IP-10. Nanofibrillated cellulose (NFC), is used here as a negative 3D gel control material due to its reported cytocompatibility and non-inflammatory properties[14]. Overall, the data indicate a limited and time-dependent inflammatory response, with no sustained broad upregulation of pro-inflammatory analytes after 72 h.

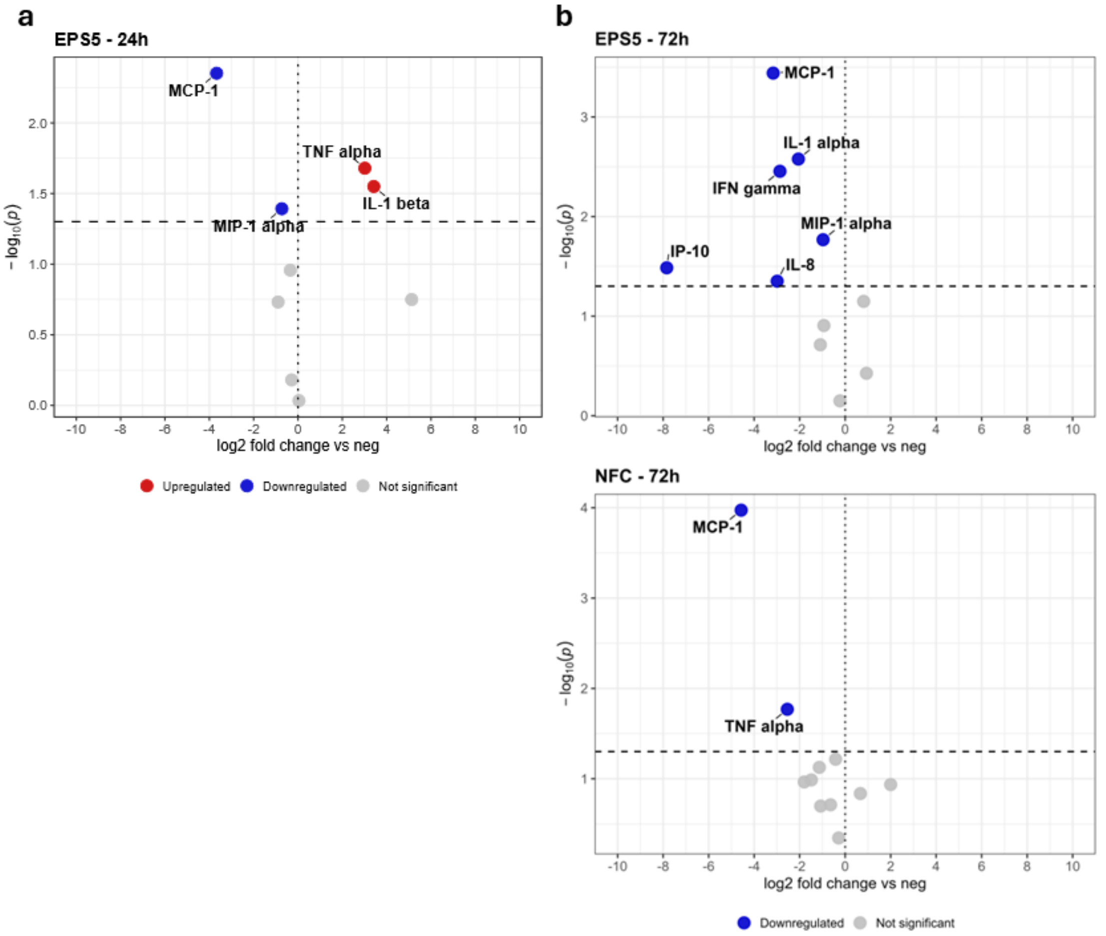


Fig. S9. Screening of immunotoxicity using THP-1-derived macrophages. Volcano plots show changes in secreted pro-inflammatory analytes relative to the negative control (unexposed macrophages) after (a) 24 h and (b) 72 h of exposure to EPS5 (pure EPS 5%) and NFC (Nanofibrillated cellulose). Red and blue points indicate significantly upregulated and downregulated analytes, respectively, while gray points indicate non-significant changes. A p-value $< 0.05$ was considered statistically significant.

**Note 5:**

Shape fidelity: The printed constructs showed a length-based shape fidelity of 91.8 ± 1.1%, indicating good agreement between the designed and printed dimensions. The filament diameter fidelity was lower and more variable (86.8 ± 11.3%), suggesting greater variability in strand width, likely due to filament spreading and small fluctuations in extrusion. Overall, the bioink maintained the intended macroscopic geometry, although further optimization of printing parameters may improve filament uniformity.

Swelling behavior: The EPSMA hydrogels showed rapid medium uptake during the initial incubation period (Fig. S10). The swelling ratio increased from approximately 35% to 175% within the first 24 h and reached approximately 215–225% after 48 h. Thereafter, the swelling ratio remained relatively constant at approximately 220–230% for up to 220 h, indicating that the hydrogels had reached swelling equilibrium. The initial increase can be attributed to diffusion of the culture medium into the hydrophilic EPSMA network. The subsequent plateau reflects a balance between medium uptake and the elastic resistance of the photocrosslinked polymer network. The absence of a marked decrease in swelling over approximately nine days suggests that the EPSMA network retained its integrity without substantial dissolution under cell-culture conditions.

Sterility: No detectable microbial growth was observed from the 5% EPS sample, while the ADP1 control showed clear growth (Fig. S11). These results indicate that the extracted EPS contained no cultivable microbial contamination under the tested conditions and was suitable for subsequent hydrogel preparation and characterization.

Degradation behavior: EPSMA hydrogels showed a rapid increase in weight during the first two days, reaching, which was attributed to swelling and media uptake. After this initial swelling phase, the control hydrogels remained stable and continued to maintain a high-water content, whereas both Proteinase K- and lysozyme-treated samples showed a gradual decrease in weight over time (Fig. S12). These results indicate that the crosslinked EPSMA hydrogel is stable in the absence of enzymes but can undergo enzyme-responsive degradation, consistent with its multicomponent composition.

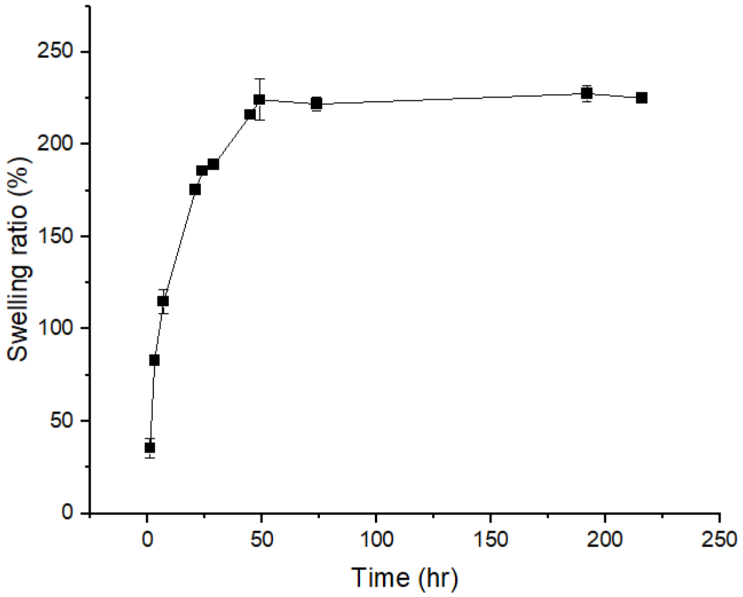


Fig. S10. Swelling behavior of EPSMA hydrogels. Photocrosslinked 5% (w/v) EPSMA hydrogels were incubated in complete 3T3 cell-culture medium at 37 °C. At predetermined time points, the samples were removed, gently blotted to remove excess surface medium, and weighed.

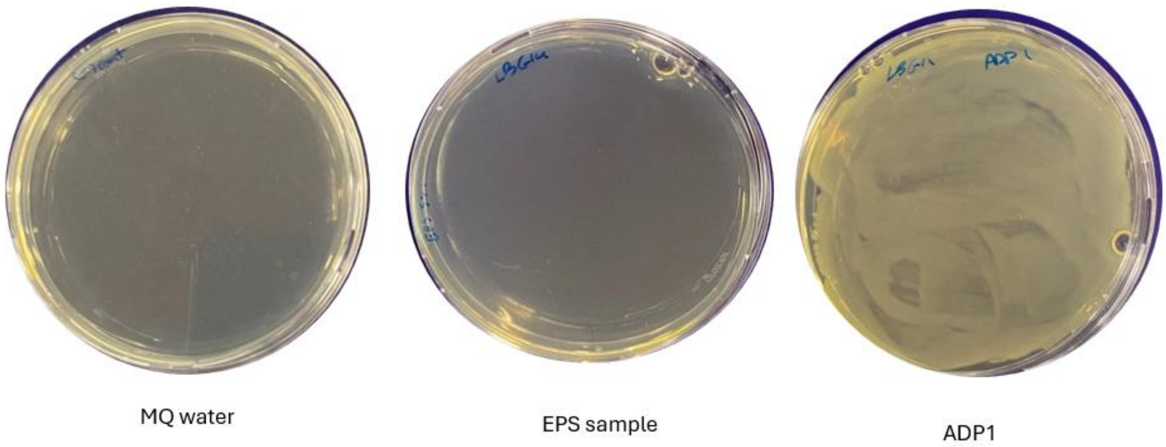


Fig. S11. Sterility assessment of the extracted EPS. 0.5% (w/v) EPS sample was spread on LA-glucose agar plates and examined for microbial growth. MQ water and *A. baylyi* ADP1 were used as negative and positive controls, respectively. No visible colonies were observed on either the MQ-water or EPS plates, whereas extensive bacterial growth was observed on the ADP1-positive-control plate.

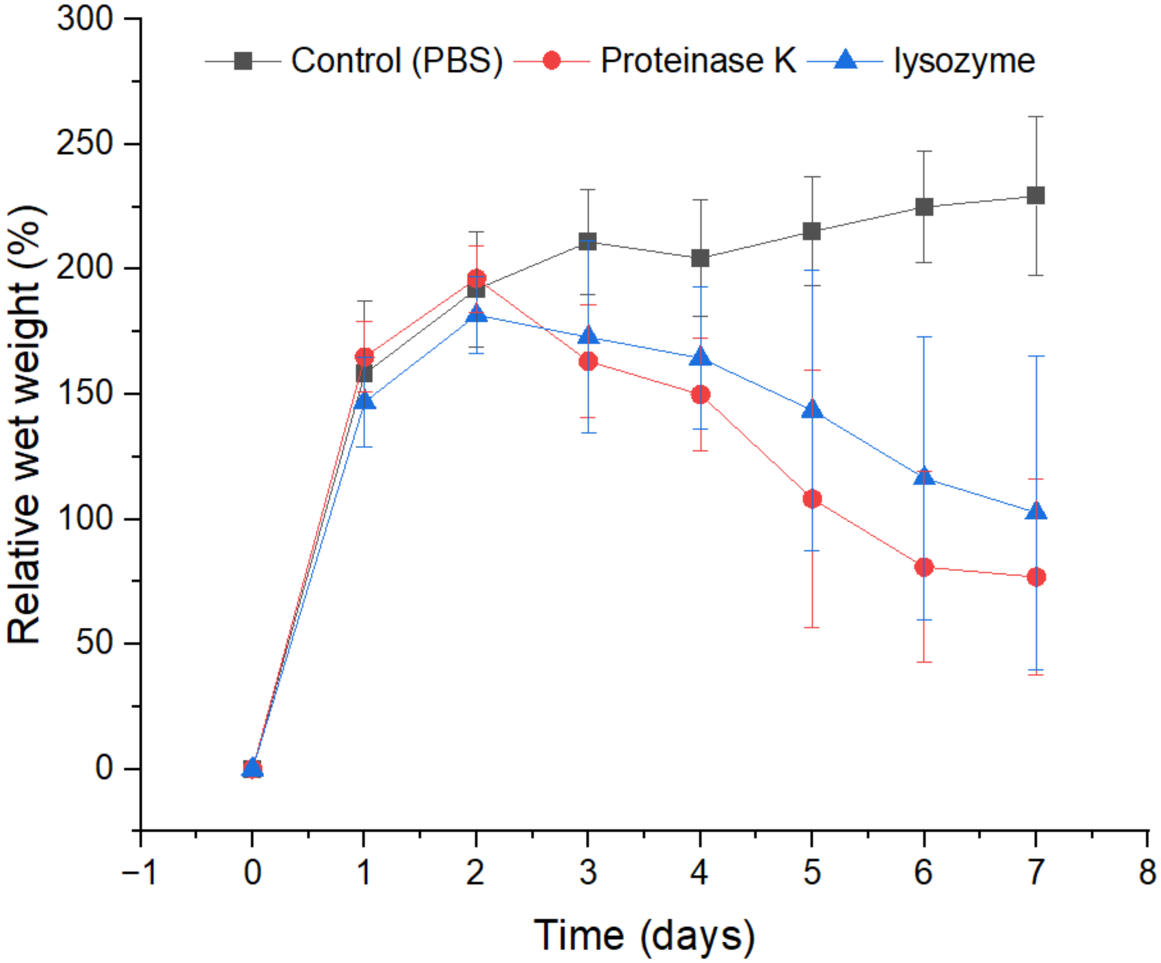


Fig. S12. Enzymatic degradation of 5% (w/v) EPSMA hydrogels in PBS without enzyme (control), with Proteinase K, and with lysozyme at 37 °C. Data are presented as mean ± SD (n = 3).

### Note 6: Cell viability

Hydrogels have great interest in biomedical applications to use as cell culture scaffolds, guiding cell growth or filling tissue deficiency[15]. One of the main requirements for hydrogels at biomedical applications is cytocompatibility and non-toxicity[16]. Here, the suitability of the EPSMA for biomedical applications was evaluated by studying cytocompatibility using widely used and robust 3T3 cells.

Among the tested concentrations, EPSMA hydrogels at 5% showed the highest cytocompatibility with over 90% cell viability (Fig. S13a, b, f), supporting good cell survival and spreading. In contrast, the 7.5% hydrogel composition resulted in slightly reduced cell density based on visual observation (Fig. S123), likely due to its higher stiffness, which may restrict cell proliferation or migration. Comparison of 2D samples with and without UV exposure shows only a slight impact on viability (Fig. S13d, e, f). Moreover, at all EPSMA gel concentrations viability was modestly higher than the 2D UV-treated control, suggesting the EPSMA hydrogel offers some protection against UV during photocrosslinking. Based on these viability results, the 5% EPSMA hydrogel was selected for subsequent 3D bioprinting experiments due to its optimal balance between printability and biocompatibility. The observed concentration-dependent cytocompatibility trend suggests that tuning EPSMA hydrogel stiffness can serve as an effective means to regulate cell–matrix interactions, which is an essential feature for designing bioinks and tissue scaffolds.

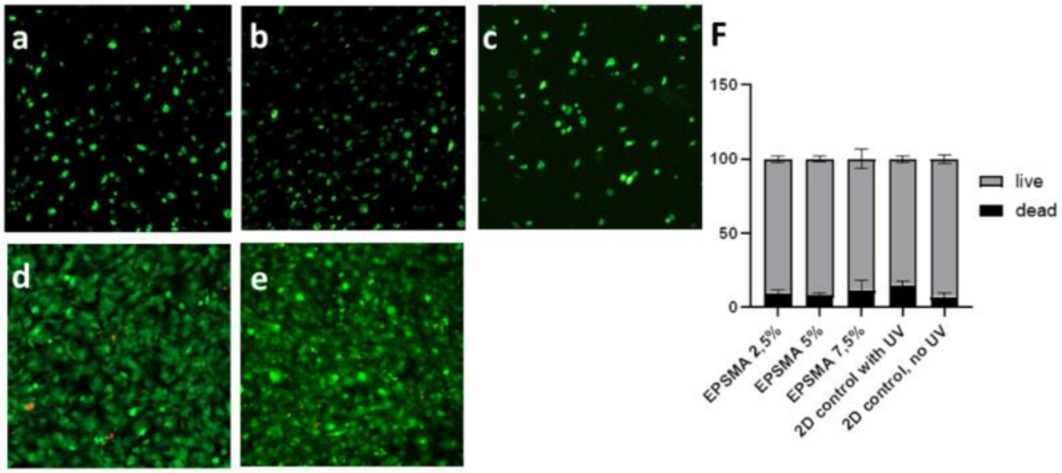


Fig. S13. Viability of the 3T3 cells inside the EPSMA hydrogels. Cells cultured encapsulated into the **a**) 2.5 %, **b**) 5%, and **c**) 7.5% EPSMA hydrogels for 6 days. Viability was analyzed using viability/cytotoxicity kit. Green = viable cells, red= dead cells. As a control, cells were cultured on 2D surface **d**) with or **e**) without UV exposure. **f**) Viable and dead cells were counted at least from 4 regions of interest per each condition and presented as percentage from all counted cells.

**Note 7:**

To evaluate whether additional cell-adhesive cues could improve cell spreading, 3T3 fibroblasts were compared in EPSMA, RGD-containing EPSMA, in which RGD (0.1 mg/mL) was physically mixed with EPSMA before crosslinking, and tissue-culture-treated plastic (TCP) as a 2D control. RGD-containing EPSMA showed a slight qualitative increase in cell elongation compared with EPSMA without RGD, although cells in both 3D hydrogel groups remained less elongated than those cultured on TCP.

The modest effect of RGD on cell elongation may be related to the mechanical stiffness of the EPSMA hydrogel. Liu et al.[17] showed that increasing hydrogel stiffness restricted 3T3 fibroblast cells elongation in alginate hydrogels, indicating that matrix mechanics can limit cell spreading despite the presence of adhesion sites. Thus, the limited elongation observed in RGD-containing EPSMA may reflect mechanical restriction imposed by the crosslinked network. Further studies should therefore explore how tuning EPSMA stiffness and network properties could enhance cell spreading and adhesion.

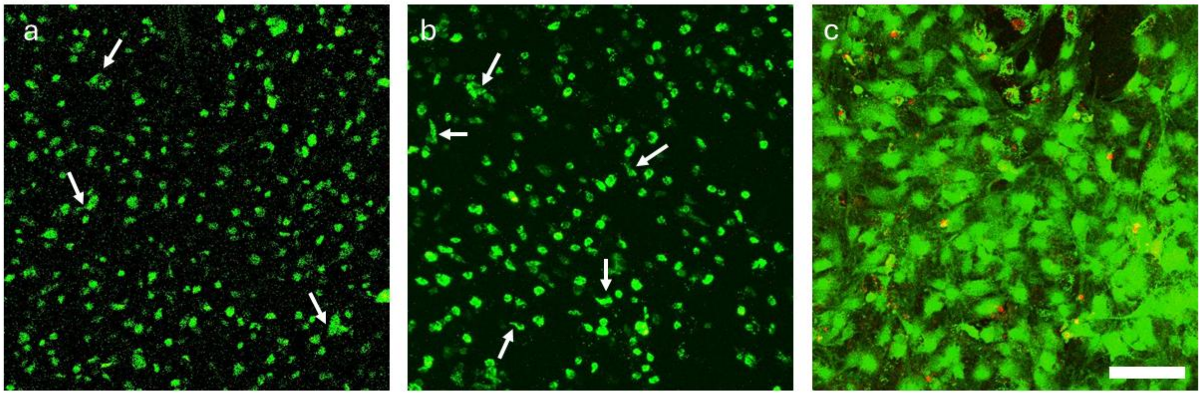


Fig. S14. Live/dead assay to 3T3 fibroblasts encapsulated into RGD-functionalized and non-functionalized EPSMA hydrogel. a) 3T3 cells grown for 6 days encapsulated inside the EPSMA hydrogel. Live/dead assay from confocal microscopy z-stack over 300 μm processed to maximum intensity projection shows good cells viability and some elongated cells (examples shown with arrows. b) 3T3 cells grown for 6 days encapsulated inside the RGD-EPSMA hydrogel. Live/dead assay from confocal microscopy z-stack over 300 μm processed to maximum intensity projection shows good cells viability and some elongated cells (examples shown with arrows). c) As a control, 2D culture of the 3T3 cells adhered to cell culture plastic. Scalebar 100 μm for all.

**DLP 3D-printer:**

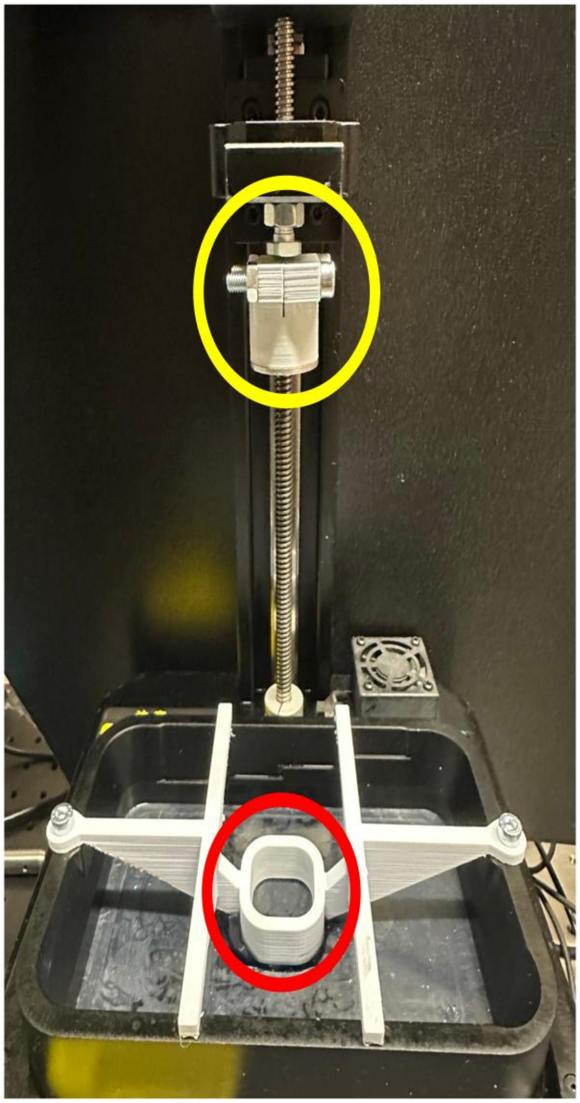

Fig. S15. Customized DLP 3D-printing setup. The yellow circle highlights a custom 2 cm build stage designed in SolidWorks; the red circle indicates a custom low-volume vat (≈4 mL capacity) for minimizing resin use. The device is a Creality LD-002R equipped with a 405 nm LED light source.

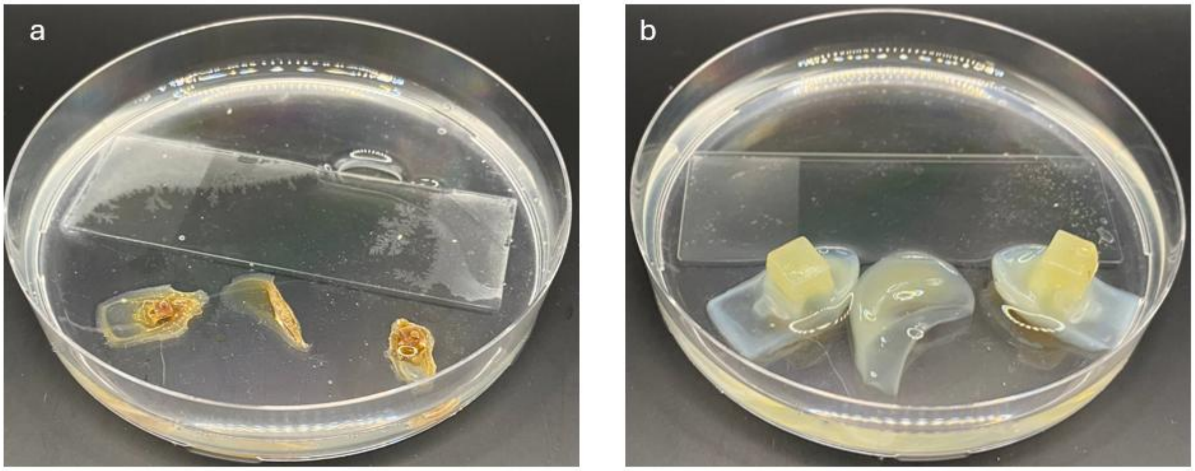


Fig. S16. After DLP printing, the EPSMA constructs were stored at room temperature for six months, and the samples became substantially dehydrated (Fig. S16a). After 12 h of immersion in water, the constructs showed substantial rehydration and largely recovered their three-dimensional geometry (Fig. S16b). These observations suggest that the crosslinked EPSMA network remained intact after prolonged dry storage and was able to recover its overall printed structure upon rehydration.

### Note 8: Injectability and in situ crosslinking

To confirm the optical behavior of the photo-initiator MB, we first measured its emission in water with Vilber Newton 7 instrument. Pure water displayed no detectable signal, whereas water containing MB showed a clear emission, confirming that the signal originates solely from MB (Fig. S17a). Next, we incorporated MB, DMA, and TEA into the 5% EPSMA hydrogel solution and monitored its response in vitro inside Eppendorf tubes. Under red-light irradiation, a time-dependent decrease in emission was observed, demonstrating the light-driven reduction of MB (Fig. S17b) and validating the instruments suitability for crosslinking detection.

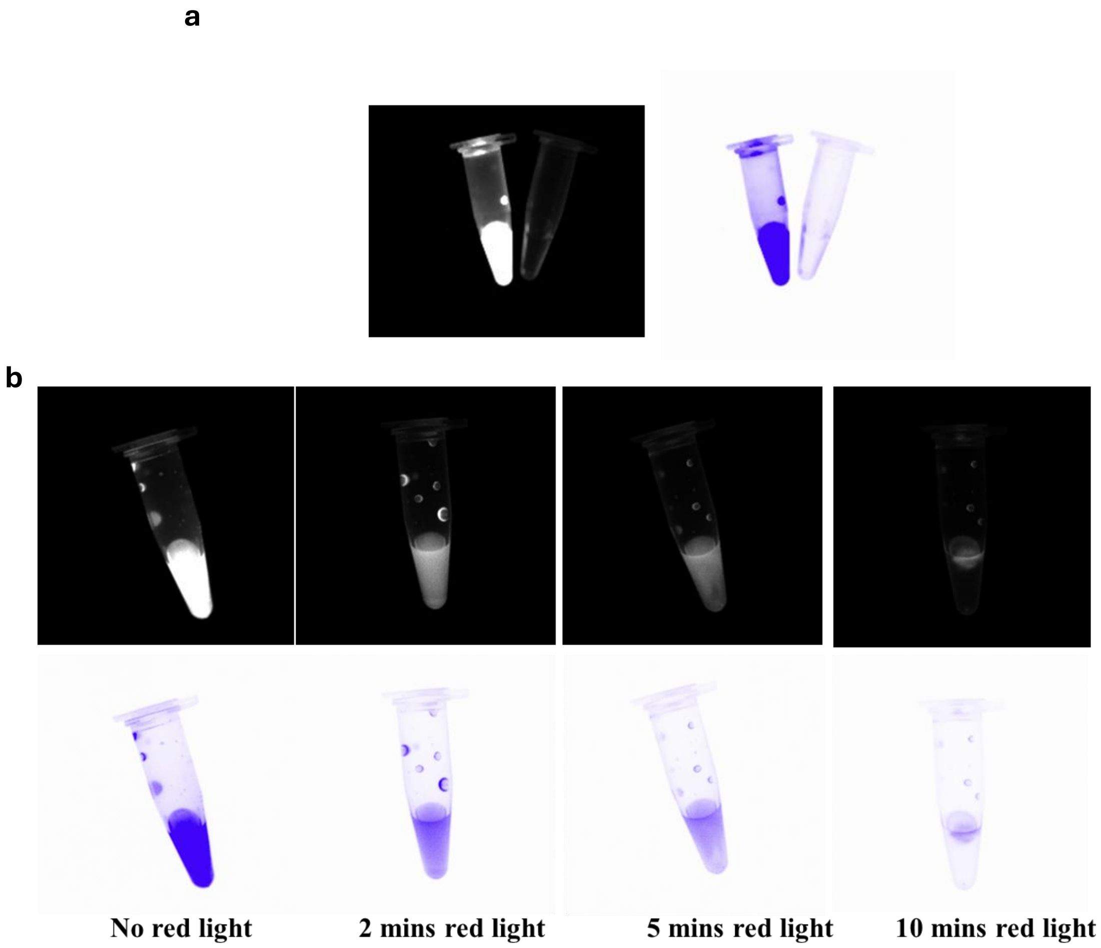


Fig. S17. Red-light crosslinking of EPSMA hydrogels monitored by Newton 7 fluorescence imaging. a) MB emission control: the MB/TEA/DMA mixture (left) shows strong fluorescence, while water (right) is dark, confirming MB-specific signal. b) Time series for a 5% (w/v) EPSMA formulation containing MB/TEA/DMA under red light for 10 min. The progressive loss of MB fluorescence indicates its photoreduction during hydrogel crosslinking.

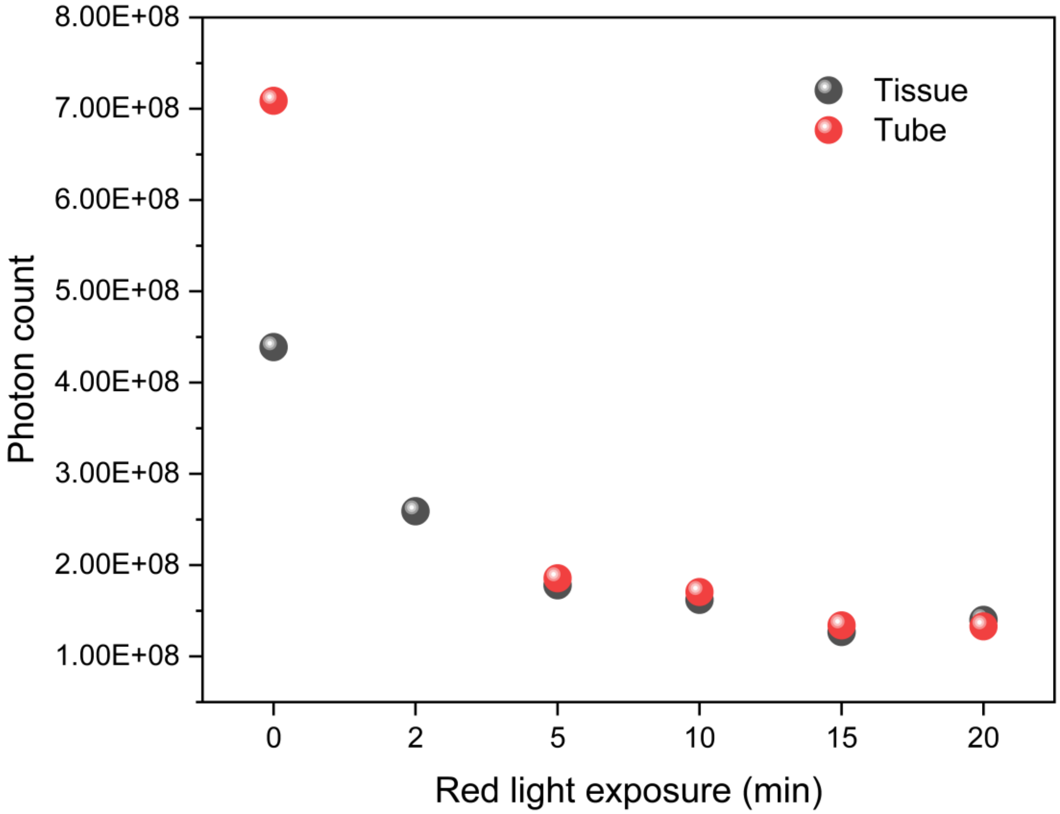


Fig. S18. Photon count by Newton 7 fluorescence imaging to monitor methylene blue (MB) emission as an indicator of crosslinking. A decrease in MB emission corresponded to hydrogel crosslinking, comparing samples crosslinked in an Eppendorf tube with those crosslinked inside chicken tissue.

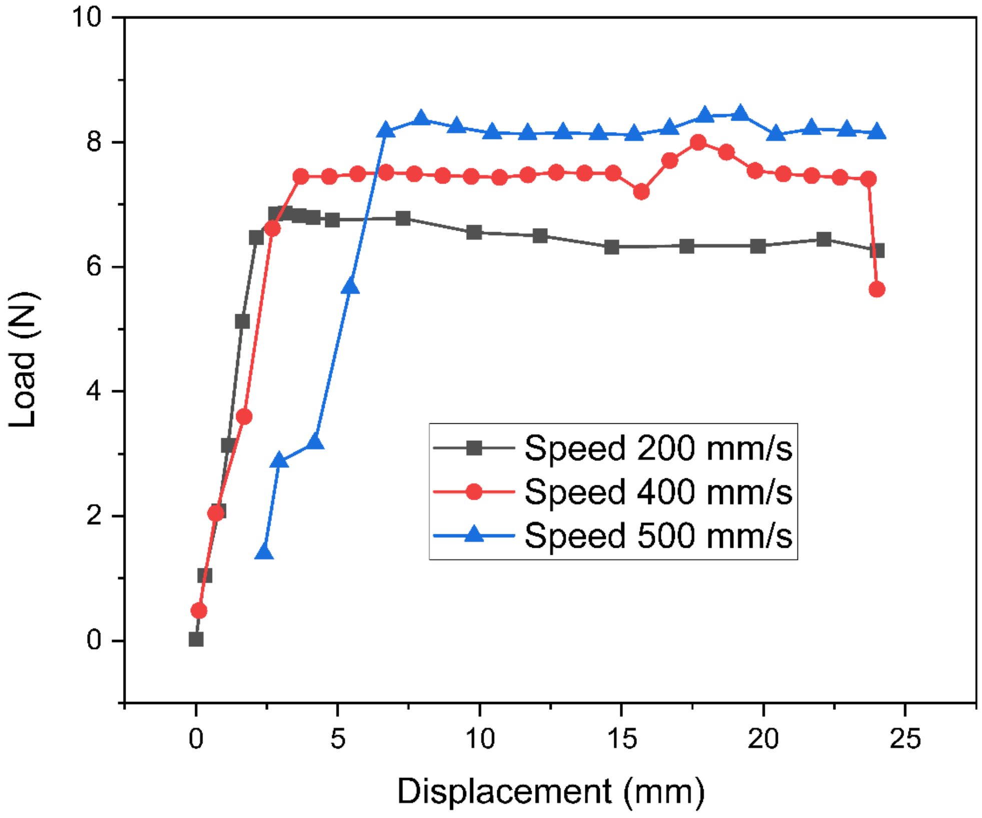


Fig. S19. Injectability test using a mechanically driven syringe. Load–displacement curves recorded while extruding the gel at three plunger speeds (200, 400, and 500 mm·s$^{-1}$). Higher speeds required greater load, with a rapid rise to a quasi-steady plateau that reflects the force needed to maintain continuous extrusion.

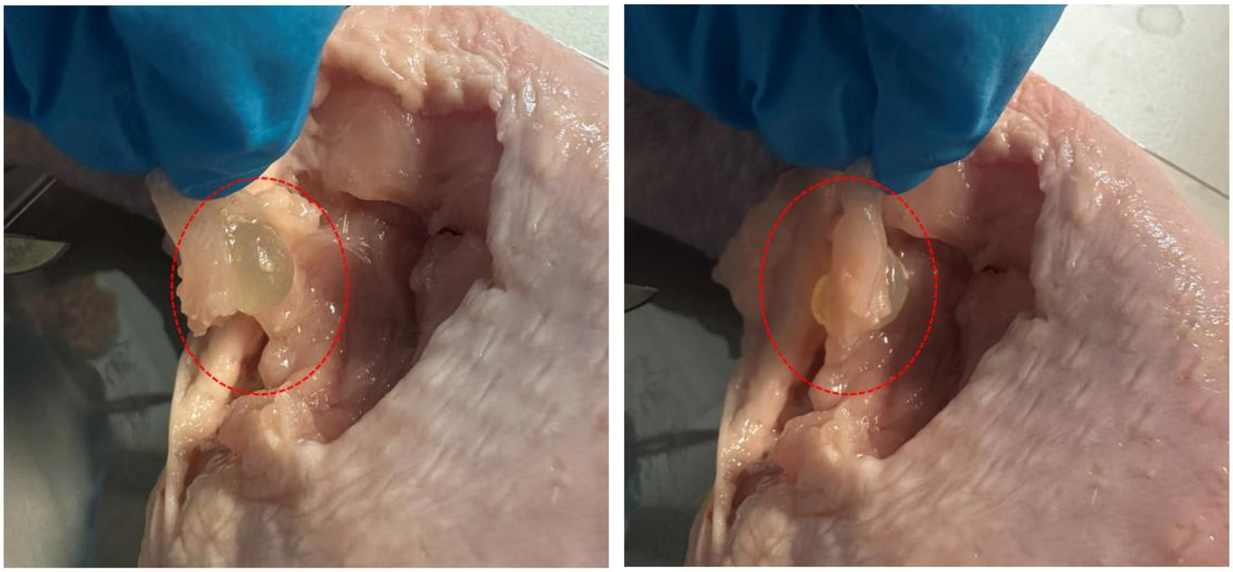

Fig. S20. Representative images of EPSMA after injection and red-light-mediated in situ crosslinking. The crosslinked gel remained localized at the injection site (red dashed circles) and was subsequently excised and weighed to determine gel recovery.

**Note 9: ABMS media composition:**

ABMS is prepared by diluting a 20× phosphate buffer solution (containing 68 g of $KH_2PO_4$ and 132.5 g of $Na_2HPO_4•H_2O$ per liter) and a 20× mineral solution (composed of 10 g of $NH_4Cl$, 5.8 g of $MgSO_4.7H_2O$, 1 g of $KNO_3$, 0.67 g of $CaCl_2•2H_2O$, 20 mg of $(NH_4)_6Mo_7O_{24}•4H_2O$, and 10 mL of SL9 solution per liter) to a 1× final concentration using water. Sodium succinate is then added to achieve a final concentration of 20 mM. SL9 is a trace metal solution made by dissolving 12.8 g of nitrilotriacetic acid, 2 g of $FeSO_4•7H_2O$, 104 mg of $CoCl_2$, 122 mg of $MnCl_2•4H_2O$, 70 mg of $ZnCl_2$, 36 mg of $Na_2MoO_4•2H_2O$, 13 mg of $NiCl_2$, 6 mg of $H_3BO_3$, and 2 mg of $CuCl_2•2H_2O$, with the pH adjusted to 6.5-7 using NaOH.

**Note 10: Materials for methacrylation and photo crosslinking:**

A dialysis membrane with a molecular weight cutoff (MWCO) of 3.5 kDa was purchased from Spectra/Por, Repligen Corp., USA. Methacrylic anhydride, trinitrobenzene sulfonic acid (TNBS), and Lithium phenyl-2,4,6-trimethylbenzoylphosphinate (LAP) were purchased from Merck KGaA, Darmstadt, Germany. DI water (deionized water, Miele Aqua Purificator G 7795, Siemens) and u.p. water (ultra-pure, Sartorius Arium Mini, 0.055 µS/cm) were used. 1H NMR analysis was carried out on an NMR spectrometer (SCZ500R, JEOL Resonance, Japan). UV-spectral measurement was performed using (Shimadzu UV-3600 plus UV−vis−NIR spectrophotometer). All chemicals and reagents used were of analytical grade. All solutions were prepared with deionized (DI) water. For red light crosslinking, Methylene blue (MB+) and N,N-Dimethylacrylamide (DMA, 99%, contains 500 ppm monomethyl ether hydroquinone as inhibitor) were purchased from Merck Darmstadt, Germany. Triethanolamine (TEA, 99+%) was purchased from Thermo Scientific Chemicals. To prepare different TEA concentrations in the millimolar (mm) range, it was diluted in deionized water.

**Table S2:** List of strains

| Strains | Relevant characteristics | Reference or source |
|---|---|---|
| ADP1 | Wild-type *A. baylyi* ADP1 | DSM 24193, DSMZ |
| ASA1400 | ADP1ΔACIAD0654::tdk/$Kan^R$ | This study |
| ASA1401 | ADP1ΔACIAD2242::tdk/$Kan^R$ | This study |
| ASA1402 | ADP1ΔACIAD2572::tdk/$Kan^R$ | This study |
| ASA1403 | ADP1ΔACIAD2279::tdk/$Kan^R$ | This study |
| ASA1404 | ADP1ΔACIAD3099::tdk/$Kan^R$ | This study |
| ASA1405 | ADP1ΔACIAD3457::tdk/$Kan^R$ | This study |
| ASA1406 | ADP1ΔACIAD3594::tdk/$Kan^R$ | This study |
| ASA1407 | ASA1400ΔACIAD0654 | This study |
| ASA1408 | ASA1407ΔACIAD2572::tdk/$Kan^R$ | This study |
| ASA1410 | ASA1409ΔACIAD3099::tdk/$Kan^R$ | This study |
| ASA1412 | ASA1411ΔACIAD3594::tdk/$Kan^R$ | This study |
| ASA1414 | ASA1413ΔACIAD2279::tdk/$Kan^R$ | This study |
| ASA1416 | ASA1415ΔACIAD3457::tdk/$Kan^R$ | This study |

**Table S3:** Primers used in this study

| ACIAD 0654 | Oligo 5'-3' |
|---|---|
| **P1** | ctctaggtgttgacgtggttg |
| **P2** | CGATGAGTTTTTCTAAGCATGCGGAGCTGGatttcaggtagttcactatgtataacaact |
| **P3** | gttagcatgactatttttaaatctccc |
| **P4** | TTTTTATGATTTGAATTGGAGGCTGGGgacaagttactccgcatttttt |
| ACIAD 2242 | Oligo 5'-3' |
| **P5** | acgcatgaccagtttttcacc |
| **P6** | CGATGAGTTTTTCTAAGCATGCGGAGCTGGtgctcgttttaatagtgaaatttaattgaact |
| **P7** | aattgaataagcacgcattaatgg |
| **P8** | TTTTTATGATTTGAATTGGAGGCTGGGctttcactttcttgatcaaaaaatcc |
| ACIAD 2572 | Oligo 5'-3' |
| **P9** | gtcactcatcaaaatggctgaattc |
| **P10** | CGATGAGTTTTTCTAAGCATGCGGAGCTGGaatgctaggcagtataaagtgaagc |
| **P11** | gaaaccagaatctggtttgc |
| **P12** | TTTTTATGATTTGAATTGGAGGCTGGGaccaatatgctatagcatttgc |
| ACIAD 2279 | Oligo 5'-3' |
| **P13** | gcttgctttggaaaaatcactctac |
| **P14** | CGATGAGTTTTTCTAAGCATGCGGAGCTGttatagattattttgttgatattcctgaataacttcaag |
| **P15** | aaaacctaaaatcgagccgaa |
| **P16** | TTTTTATGATTTGAATTGGAGGCTGGGctgctttataagactatgtgtagg |
| ACIAD 3099 | Oligo 5'-3' |
| **P17** | ctaaaggccatgtaattggcac |
| **P18** | CGATGAGTTTTTCTAAGCATGCGGAGCTGGgttagatttaccatcaattggtcatataacc |
| **P19** | agtcacaactttaccgtgatg |
| **P20** | TTTTTATGATTTGAATTGGAGGCTGGGaaaggacttccatgcgtattt |
| ACIAD 3457 | Oligo 5'-3' |
| **P21** | aagaatctgtaaagctgggcttag |
| **P22** | CGATGAGTTTTTCTAAGCATGCGGAGCTGGcttgatgagaggcatttatttaaaataaaatttcac |
| **P23** | ctggggtcattgtattgtaaca |
| **P24** | TTTTTATGATTTGAATTGGAGGCTGGGttaaaattagaataggctgtttctgtc |
| ACIAD 3594 | Oligo 5'-3' |
| **P25** | gcggacaaaacctttttatgtaacc |
| **P26** | CGATGAGTTTTTCTAAGCATGCGGAGCTGtgcttaatccagttcttcattgacttg |
| **P27** | gatcaaaggtggagaagatcc |

| | |
|---|---|
| **P28** | TTTTTATGATTTGAATTGGAGGCTGGGcgcacttattcaatatgaattttttatatc |
| | (**Rescue primers**) Oligo 5'-3' |
| **P29** | aaaaaatgcggagtaacttgtcatttcaggtagttcactatgtataacaact |
| **P30** | catagtgaactacctgaaatgacaagttactccgcatttttt |
| **P31** | tttgatcaagaaagtgaaagtgctcgttttaatagtgaaatttaattgaact |
| **P32** | aaatttcactattaaaacgagcactttcactttcttgatcaaaaaatcc |
| **P33** | aaatgctatagcatattggtaatgctaggcagtataaagtgaagc |
| **P34** | actttatactgcctagcattaccaatatgctatagcatttgc |
| **P35** | aaatacgcatggaagtcctttgttagatttaccatcaattggtcatataacc |
| **P36** | caattgatggtaaatctaacaaaggacttccatgcgtattt |
| **P37** | aaaaaaattcatattgaataagtgcgtgcttaatccagttcttcattgacttg |
| **P38** | aatgaagaactggattaagcacgcacttattcaatatgaattttttatatc |
| **P39** | cctacacatagtcttataaagcagttatagattattttgttgatattcctgaataacttcaag |
| **P40** | ggaatatcaacaaaataatctataactgctttataagactatgtgtagg |

Red: overlap with tdk-kan cassette.
